# WNK-Dependent Phosphorylation of Gephyrin Tunes GABA_A_ Receptors at Inhibitory Synapses and Modulates Anxiety Behavior

**DOI:** 10.1101/2025.09.05.674425

**Authors:** Zaha Merlaud, Celia Delhaye, Margarida Nabais, Zahra Imani, Erwan Pol, Marion Russeau, Maelys Tostain, Juliette Gouhier, Romane Rahir, Corentin Le Magueresse, Marika Norsten-Bertrand, Sabine Lévi

## Abstract

The role of the chloride-sensitive kinase WNK1 and its effector SPAK in the brain remains poorly understood. Here, we identify a regulatory mechanism involving WNK signaling that directly controls the synaptic diffusion and clustering, as well as the membrane stability and endocytosis of inhibitory GABA_A_ receptors (GABA_A_Rs). We show that activation of WNK signaling stabilizes GABA_A_Rs at inhibitory synapses, while inhibition enhances receptor internalization. This regulation depends on the phosphorylation state of two previously uncharacterized residues in the central linker region of the gephyrin scaffold protein. Modulating WNK activity alters neuronal activity and the kinetics of GABAergic currents. *In vivo*, expression of a phospho-mimetic form of gephyrin at WNK-targeted sites produces anxiolytic effects. By orchestrating the recruitment of GABA_A_Rs at inhibitory synapses, the WNK pathway emerges as a master regulator of GABAergic transmission and establishes chloride as a bona fide second messenger in inhibitory synaptic signaling.

**Significance:** Synaptic transmission relies on signaling pathways that control how neurotransmitter receptors move and stabilize at synapses. Many of these pathways use calcium as a second messenger. In contrast, how chloride might regulate synapses has remained unclear. In this study, we identify a chloride-sensitive pathway involving the WNK1 kinase and its partner SPAK, which controls the movement and stability of inhibitory GABA_A_ receptors at synapses. This occurs through the phosphorylation of gephyrin, a key scaffolding protein, at two newly identified sites. Our results show that intracellular chloride can act as a second messenger, reshaping inhibitory synapses and altering behavior in mice, including reducing anxiety-like responses.

## Introduction

The chloride-permeable ionotropic γ-aminobutyric acid receptor (GABA_A_R) is the primary neurotransmitter receptor mediating inhibition in the brain. The strength of GABAergic synaptic transmission depends critically on the number and spatial organization of GABA_A_Rs at the synapse. A key mechanism regulating receptor availability is the diffusion-capture process, which slows and traps receptors at the postsynaptic membrane, leading to their accumulation at inhibitory synapses (Choquet and Triller, 2003; Triller and Choquet, 2005; Choquet and Triller, 2013; Jacob et al., 2005; Mukherjee et al., 2011; Bannai et al., 2009). This process is controlled by the interaction of GABA_A_Rs with gephyrin, the main scaffolding protein that forms a dynamic submembrane network at inhibitory sites. Activity-dependent modulation of receptor diffusion and clustering is influenced by post-translational modifications of both GABA_A_Rs and gephyrin, particularly phosphorylation, which underlies synaptic plasticity (Bannai et al., 2009, 2015; Muir et al., 2010; Petrini et al., 2014; de Luca et al., 2017; Ravasenga et al., 2022; Chiu et al., 2019; Barberis, 2020; Battaglia et al., 2018; Flores et al., 2015; Petrini et al., 2014; Tyagarajan et al., 2011, 2013).

The chloride-sensitive WNK1 kinase pathway, known to regulate chloride transporter activity *via* its effectors SPAK and OSR1, is implicated in neurological disorders with inhibitory dysfunction such as epilepsy and neuropsychiatric diseases (references in Shekarabi et al., 2017; McMoneagle et al., 2024). WNK1 activation under low-chloride conditions (Piala et al., 2014) phosphorylates KCC2 at threonine residues T906/T1007 and NKCC1 at T203/T207/T212 (Inoue et al., 2012; Thastrup et al., 2012; de los Heros et al., 2014; Friedel et al., 2015), modulating their membrane dynamics, clustering, and stability, which alters intracellular chloride concentration (Heubl et al., 2017; Côme et al., 2023). However, whether WNK1 signaling acts directly at inhibitory synapses to regulate GABA_A_Rs and gephyrin is unknown.

We show that acute activation of the WNK pathway rapidly modulates membrane dynamics of the synaptic GABA_A_R α1 subunit (GABA_A_R α1), promoting its confinement and clustering at inhibitory synapses alongside gephyrin recruitment. Conversely, WNK inhibition reduces GABA_A_R α1 clustering and enhances receptor internalization. This synaptic reorganization depends on the phosphorylation state of two previously uncharacterized residues, threonine T260 and serine S280, in the gephyrin linker region. Finally, intrahippocampal expression of phosphomimetic T260E/S280E-gephyrin increases synaptic GABA_A_R α1 levels *in vivo*, alters the kinetics of GABAergic currents, and reduces anxiety in mice. These findings reveal a novel WNK-dependent mechanism regulating synaptic inhibition with potential therapeutic relevance in anxiety disorders.

## Results

### Inhibition of WNK signaling promotes GABA_A_R endocytosis

We examined the role of the WNK pathway in regulating GABA_A_Rs containing the α1 subunit, which is enriched at inhibitory synapses in the hippocampus (Fritschy and Mohler, 1995; Pirker et al., 2000; Sieghart and Sperk, 2002; Brünig et al., 2002). To this end, we analyzed the impact of WNK signaling on the membrane dynamics of GABA_A_R α1 using quantum dot-based single-particle tracking (QD-SPT) in primary hippocampal cultures exhibiting functional GABAergic inhibition (Heubl et al., 2017). Given that GABA_A_R lateral diffusion is rapidly modulated by glutamatergic activity (Bannai et al., 2015, 2009; Battaglia et al., 2018; Muir et al., 2010; Petrini et al., 2014), experiments were performed in the presence of tetrodotoxin (TTX, 1 μM), kynurenic acid (KYN, 1 mM), and the group I/II mGluR antagonist R,S-MCPG (500 μM) to silence glutamatergic transmission. To acutely inhibit the WNK pathway, neurons were treated with either the SPAK/OSR1 inhibitor closantel or the pan-WNK inhibitor WNK463 prior to QD-SPT experiments. We found that GABA_A_R α1 explored a smaller area of the extrasynaptic membrane in neurons treated with closantel or WNK463 compared to control conditions (**Figure S1A**). In contrast, at inhibitory synapses, GABA_A_R α1 trajectories were unaffected by drug treatment **(Figure S1A**). Consistent with these findings, quantitative analyses revealed that, outside synapses, the receptor exhibited decreased mobility and increased confinement **(Figure S1B-C),** whereas the diffusion coefficient **(Figure S1B),** explored area **(Figure S1C),** and dwell time at GABAergic synapses **(Figure S1D)** remained unchanged following WNK/SPAK pathway inhibition.

GABA_A_Rs have been reported to exhibit reduced lateral diffusion within clathrin-enriched endocytic zones (EZs), which are predominantly localized in extrasynaptic regions (Smith et al., 2012; Merlaud et al., 2022). To investigate whether WNK signaling promotes the targeting of GABA_A_Rs to endocytic zones, we disrupted EZ formation using a peptide (DIP, dynamin inhibitory peptide) that interferes with the dynamin–amphiphysin interaction. As shown in **Figure S1A**, when endocytosis is blocked with DIP, the extrasynaptic trajectories of GABA_A_R α1 in the presence of WNK or SPAK inhibitors are indistinguishable from those observed in the control + DIP condition. Quantitative analyses confirmed these findings. Specifically, acute exposure to WNK463 or closantel in the presence of DIP no longer affected the diffusion coefficient or confinement of GABA_A_R α1 in the extrasynaptic membrane (**Figure S1 E-F**). Similarly, blocking endocytosis did not alter GABA_A_R α1 dynamics at synapses. The diffusion coefficient, explored area, and dwell time at inhibitory synapses remained unchanged following acute treatment with WNK or SPAK inhibitors (**Figure S1E, S1F, and S1G**, respectively). **Accordingly, our data indicate that under basal activity conditions, acute inhibition of WNK signaling restricts GABA_A_R α1 to extrasynaptic endocytic zones, without affecting synaptic receptor dynamics.**

To determine whether the increased confinement of GABA_A_R α1 to endocytic zones following acute WNK/SPAK pathway inhibition is associated with enhanced internalization, we quantified the surface expression of GABA_A_R α1 in the presence or absence of the WNK inhibitor. We calculated the ratio of surface to total (surface + intracellular) GABA_A_R α1 fluorescence intensity **(Figure S2A)** and observed a significant decrease in this ratio after a 1-hour exposure to WNK463 **(Figure S2B). These results indicate that GABA_A_R α1 undergoes internalization upon acute inhibition of WNK signaling.**

We next investigated whether blocking WNK activity alters the surface clustering of GABA_A_R α1 using immunocytochemistry on non-permeabilized cells. Given that the size of GABA_A_R α1 clusters is at the limit of resolution for a standard epifluorescence microscope, we further analyzed the impact of treatments on clustering using Stochastic Optical Reconstruction Microscopy (STORM). STORM images revealed an overall reduction in GABA_A_R α1 membrane clustering **(Figure S2C),** which was corroborated by quantitative analysis showing that acute treatments with closantel and WNK463 significantly decreased both the size of GABA_A_R α1 clusters and their single molecule detection density **(Figure S2D).** This reduction in clustering, combined with the decreased surface expression of GABA_A_Rs **(Figure S2A-B)** and the increased receptor confinement within endocytic zones **(Figure S1),** supports the hypothesis that **WNK blockade leads to the dislocation of GABA_A_R clusters, followed by receptor recruitment into endocytic zones and subsequent internalization.**

### Activation of the WNK pathway stabilizes GABA_A_Rs and the gephyrin scaffold at inhibitory synapses

Given that blockade of WNK or SPAK kinases promotes GABA_A_R α1 internalization, we hypothesized that stimulating WNK signaling would, in contrast, stabilize GABA_A_Rs at the membrane. Since no direct pharmacological activators of WNK or SPAK have been developed, we activated the WNK pathway by using an extracellular solution depleted of chloride (2 mM [Cl^−^]). This solution reduces intracellular Cl^−^ concentration, promoting WNK autophosphorylation and the regulation of KCC2 and NKCC1 membrane diffusion and clustering (Moriguchi et al., 2005; Piala et al., 2014; Heubl et al., 2017; Côme et al., 2023). We then assessed the lateral diffusion of GABA_A_R α1 under these conditions **(Figure 1).** In the extrasynaptic compartment, we observed increased surface exploration of individual quantum dots (QDs) **(Figure 1A)** as well as for a larger population of QDs **(Figure 1C),** indicating reduced confinement of GABA_A_R α1 in low versus high chloride conditions. This was accompanied by an increase in the diffusion coefficient for extrasynaptic GABA_A_R α1 **(Figure 1A-B),** suggesting that WNK activation releases diffusion constraints for these receptors. The addition of closantel or WNK463 to the chloride-depleted solution prevented these effects **(Figure 1A-C),** demonstrating the involvement of the WNK pathway in this regulation. Furthermore, while acute WNK blockade at steady state had no effect on GABA_A_R α1 mobility at inhibitory synapses **(Figure S1),** activation of the WNK pathway reduced GABA_A_R α1 surface exploration **(Figure 1C)** and increased the receptor’s dwell time at the synapse **(Figure 1D).** Thus, activation of the WNK pathway confines GABA_A_R α1 to the GABAergic synapse. Interestingly, while WNK463 in the 2 mM [Cl^−^] solution prevented the increased receptor confinement elicited by low chloride conditions at extrasynaptic sites **(Figure 1B-D),** closantel unexpectedly enhanced the effects of WNK pathway activation on GABA_A_R α1 diffusion coefficient **(Figure 1B),** explored area **(Figure 1C),** and dwell time at the synapse **(Figure 1D).** Given the reduced clustering of GABA_A_R α1 upon closantel treatment at steady state **(Figure S2D),** we propose that this increased confinement results from receptor trapping and internalization in perisynaptic endocytic zones. Altogether, these findings reveal that **WNK pathway activation increases the trapping of GABA_A_Rs at GABAergic synapses.**

**Figure 1.**
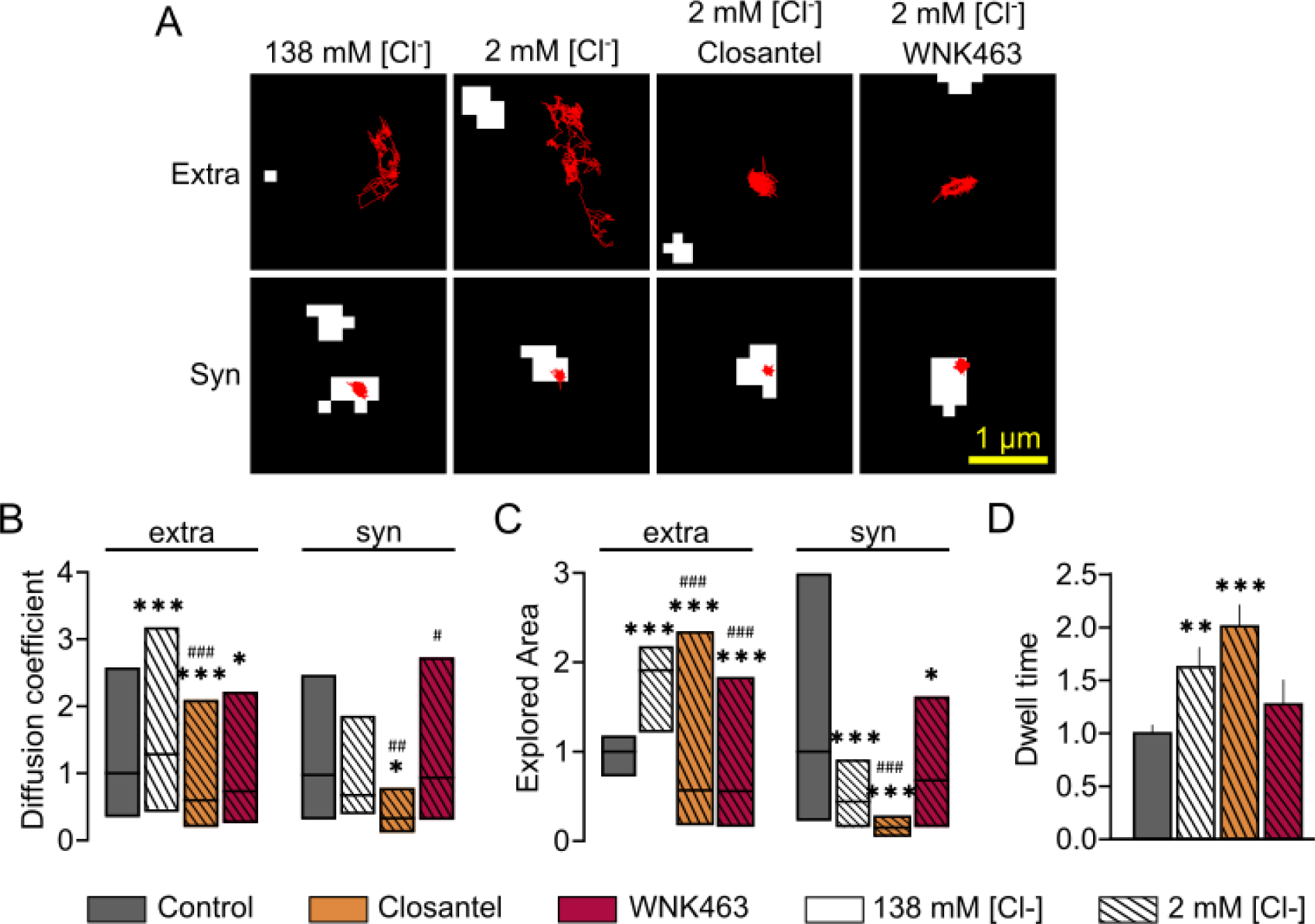
The WNK pathway tunes GABA_A_R α1 lateral diffusion in hippocampal cultured neurons. **A.** Examples of extrasynaptic (bottom) or synaptic (top) GABA_A_R α1 reconstructed trajectories (red) for neurons treated with 138 mM [Cl-], 2 mM [Cl-], 2 mM [Cl-] + closantel or 2 mM [Cl-] + WNK463. White spots represent gephyrin clusters. **B-C.** Diffusion coefficient (B) and explored area (C) for extrasynaptic (extra) and synaptic (syn) GABA_A_R α1 for neurons treated with 2 mM [Cl-] (hatched white), 2 mM [Cl-] + closantel (hatched orange), 2 mM [Cl-] + WNK463 (hatched red), as compared to neurons exposed to 138 mM [Cl-] (plain grey). ***B:*** 138 mM [Cl-] extra n=1335 QDs, syn n=206 QDs, 2 mM [Cl-] extra n=1418 QDs, syn n=118 QDs, 2 mM [Cl-] + closantel extra n=687 QDs, syn n=66 QDs and 2 mM [Cl-] + WNK463 extra n=426 QDs, syn n=66 QDs, 3 cultures. Extra: 2 mM [Cl-] vs 138 mM [Cl-] p<0.0001, 2 mM [Cl-] + closantel vs 138 mM [Cl-] p<0.0001, vs 2 mM [Cl-] p<0.0001, 2 mM [Cl-] + WNK463 vs 138 mM [Cl-] p=0.0319, vs 2 mM [Cl-] p<0.0001; syn: 2 mM [Cl-] vs 138 mM [Cl-] p=0.1407, 2 mM [Cl-] + closantel vs 138 mM [Cl-] p=0.0001, vs 2 mM [Cl-] p=0.0001, 2 mM [Cl-] + WNK463 vs 138 mM [Cl-] p=0.8862, vs 2 mM [Cl-] p=0.2917; ***C:*** 138 mM [Cl-] extra n=4400 QDs, syn n=351 QDs, 2 mM [Cl-] extra n=4498 QDs, syn n=128 QDs, 2 mM [Cl-] + closantel extra n=2152 QDs, syn n=109 QDs and 2 mM [Cl-] + WNK463 extra n=579 QDs, syn n=97 QDs, 3 cultures. Extra: 2mM [Cl-] vs 138 mM [Cl-] p<0.0001, 2 mM [Cl-] + closantel vs 138 mM [Cl-] p<0.0001, vs 2 mM [Cl-] p<0.0001, 2 mM [Cl-] + WNK463 vs 138 mM [Cl-] p<0.0001, vs 2 mM [Cl-] p<0.0001; syn: 2 mM [Cl-] vs 138 mM [Cl-] p<0.0001, 2 mM [Cl-] + closantel vs 138 mM [Cl-] p<0.0001, vs 2 mM [Cl-] p<0.0001, 2 mM [Cl-] + WNK463 vs 138 mM [Cl-] p=0.0272, vs 2 mM [Cl-] p=0.1191. Data are presented as median values ± 25–75% IQR. Values were normalized and compared to the corresponding control values: * vs 138 mM [Cl-], # vs 2 mM [Cl-]. Kolmogorov-Smirnov test. **D.** Dwell time of GABA_A_R α1 for neurons treated with 138 mM [Cl-] (plain grey), 2 mM [Cl-] (hatched white), 2 mM [Cl-] + closantel (hatched orange) or 2 mM [Cl-] + WNK463 (hatched red). D: 138 mM [Cl-] n=410 QDs, 2 mM [Cl-] n=191 QDs, 2 mM [Cl-] + closantel n=214 QDs and 2 mM [Cl-] + WNK463 n=92 QDs, 3 cultures. 2 mM [Cl-] vs 138 mM [Cl-] p=0.0042, 2 mM [Cl-] + closantel vs 138 mM [Cl-] p<0.0001, vs 2 mM [Cl-] p>0.9999, 2 mM [Cl-] + WNK463 vs 138 mM [Cl-] p>0.9999, vs 2 mM [Cl-] p=0.4761. Data are presented as mean values ± SEM. Values were normalized and compared to the corresponding control values: * vs 138 mM [Cl-], # vs 2 mM [Cl-]. Kruskal-Wallis test.

We thus hypothesized that the increased confinement of GABA_A_R α1 at synapses would lead to receptor accumulation at these sites. We assessed the impact of WNK activation on the surface pool of GABA_A_R α1. In contrast to the loss of GABA_A_R α1 membrane expression induced by WNK blockade **(Figure S2A-B),** we observed that WNK stimulation increased the surface-to-total GABA_A_R α1 ratio **(Figure 2A-B),** indicating **enhanced membrane stability of GABA_A_R α1.** Furthermore, this increase in the surface pool of GABA_A_R α1 was prevented by the addition of WNK463 to the chloride-depleted solution **(Figure 2A-B),** demonstrating the involvement of the WNK pathway in this effect.

**Figure 2.**
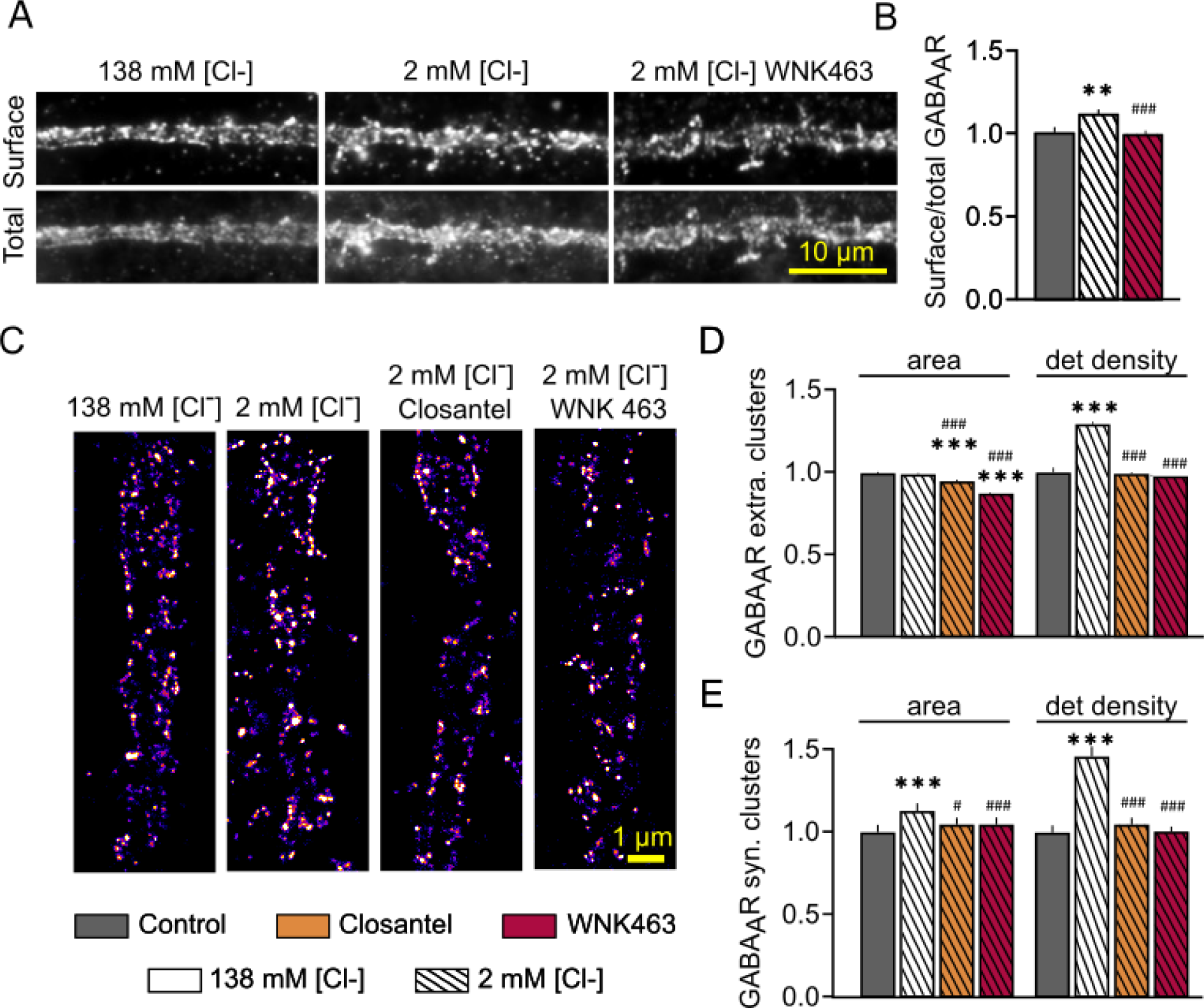
Activation of the WNK pathway increases GABA_A_R α1 at inhibitory synapses. **A.** Surface (top) and total (surface + intracellular) GABA_A_R α1 (bottom) immunostaining in neurons exposed to 138 mM [Cl-] (plain grey), 2 mM [Cl-] (hatched white) or 2 mM [Cl-] (hatched red). **B.** Quantification of the surface/total ratio of GABA_A_R α1. Control n=58 cells, 2 mM [Cl-] n=61 cells, 2 mM [Cl-] + WNK463 n=59 cells, 3 cultures. 2 mM [Cl-] vs 138 mM [Cl-] p=0.0012, 2 mM [Cl-] + WNK463 vs 138 mM [Cl-] p>0.9999, vs 2 mM [Cl-] p=0.0004. **C.** STORM rendered images of GABA_A_R α1 for 138 mM [Cl-], 2 mM [Cl-], 2 mM [Cl-] + closantel and 2 mM [Cl-] + WNK463 treated neurons. Warmer colors correspond to higher detection density. **D-E.** Quantification of the mean surface area and detection density of GABA_A_R α1 extrasynaptic (D) and synaptic clusters (E) for neurons treated with 138 mM [Cl-] (plain grey), 2 mM [Cl-] (hatched white), 2 mM [Cl-] + closantel (hatched orange) or 2 mM [Cl-] + WNK463 (hatched red). ***D:*** 138 mM [Cl-] n=3258 clusters, 2mM [Cl-] n=2723 clusters, 2mM [Cl-] + closantel n=2300 clusters, 2mM [Cl-] + WNK463 n=2395 clusters, 3 cultures. Area: 2mM [Cl-] vs 138 mM [Cl-] p>0.9999, 2 mM [Cl-] + closantel vs 138 mM [Cl-] p<0.0001 vs 2 mM [Cl-] p<0.0001, 2 mM [Cl-] + WNK463 vs 138 mM [Cl-] p<0.0001, vs 2 mM [Cl-] p<0.0001; detection density: 2mM [Cl-] vs 138 mM [Cl-] p<0.0001, 2 mM [Cl-] + closantel vs 138 mM [Cl-] p>0.9999, vs 2 mM [Cl-] p<0.0001, 2 mM [Cl-] + WNK463 vs 138 mM [Cl-] p=0.2502, vs 2 mM [Cl-] p<0.0001; ***E:*** 138 mM [Cl-] n=1145 clusters, 2mM [Cl-] n=1071 clusters, 2mM [Cl-] + closantel n=974 clusters, 2mM [Cl-] + WNK463 n=789 clusters, 3 cultures. Area: 2mM [Cl-] vs 138 mM [Cl-] p<0.0001, 2 mM [Cl-] + closantel vs 138 mM [Cl-] p=0.2584 vs 2 mM [Cl-] p=0.0416, 2 mM [Cl-] + WNK463 vs 138 mM [Cl-] p>0.9999, vs 2 mM [Cl-] p=0.0002; detection density: 2mM [Cl-] vs 138 mM [Cl-] p<0.0001, 2 mM [Cl-] + closantel vs 138 mM [Cl-] p>0.9999, vs 2 mM [Cl-] p<0.0001, 2 mM [Cl-] + WNK463 vs 138 mM [Cl-] p=0.0971, vs 2 mM [Cl-] p<0.0001. Data are presented as mean values ± SEM. Values were normalized and compared to the corresponding control values: * vs 138 mM [Cl-], # vs 2 mM [Cl-]. Kruskal-Wallis test.

We then investigated whether activation of the WNK pathway could modulate the clustering of synaptic and extrasynaptic GABA_A_R α1 using STORM. Clusters were categorized based on size, with the largest considered synaptic, as previously described (Merlaud et al., 2022). Activation of the WNK pathway using low chloride solution initially increased GABA_A_R α1 clustering, regardless of cluster size **(Figure 2C),** an effect that was abolished in the presence of WNK or SPAK inhibitors **(Figure 2C).** Quantification confirmed these observations. Small ‘extrasynaptic’ clusters showed increased detection density upon WNK pathway activation, without changes in their overall surface area **(Figure 2D).** This effect was blocked by the addition of WNK or SPAK inhibitors to the low chloride solution **(Figure 2C-D),** suggesting an increase in receptor content within extrasynaptic membrane clusters. Moreover, activation of the WNK pathway significantly increased both the area and detection density of large ‘synaptic’ clusters **(Figure 2 C, E),** implying receptor recruitment to synapses, which was prevented by the addition of pathway inhibitors **(Figure 2 C, E).** Altogether, these findings indicate that activation of **the WNK pathway stabilizes GABA_A_Rs within membrane clusters at both synaptic and extrasynaptic sites.**

Synaptic GABA_A_Rs interact with the key scaffolding molecule gephyrin, which tightly regulates the lateral diffusion and clustering of the receptors through a diffusion-trapping mechanism (Petrini and Barberis 2014). In this context, we investigated the effect of WNK signaling on gephyrin clustering at synapses. Using conventional fluorescence microscopy, we quantified gephyrin clusters colocalized with the vesicular GABA transporter VGAT **(Figure 3A)** following blockade of either WNK or SPAK kinases. Overall, blockade of the pathway did not alter gephyrin clustering at extrasynaptic sites **(Figure 3B).** However, it significantly reduced gephyrin clustering at GABAergic synapses **(Figure 3C).** Consistent with the reduced synaptic clustering of GABA_A_R α1 upon WNK or SPAK inhibition **(Figure S2),** this result suggests that **the decreased receptor density at synapses under WNK/SPAK blockade conditions is likely due to a reduction in the availability of scaffolding proteins.**

**Figure 3.**
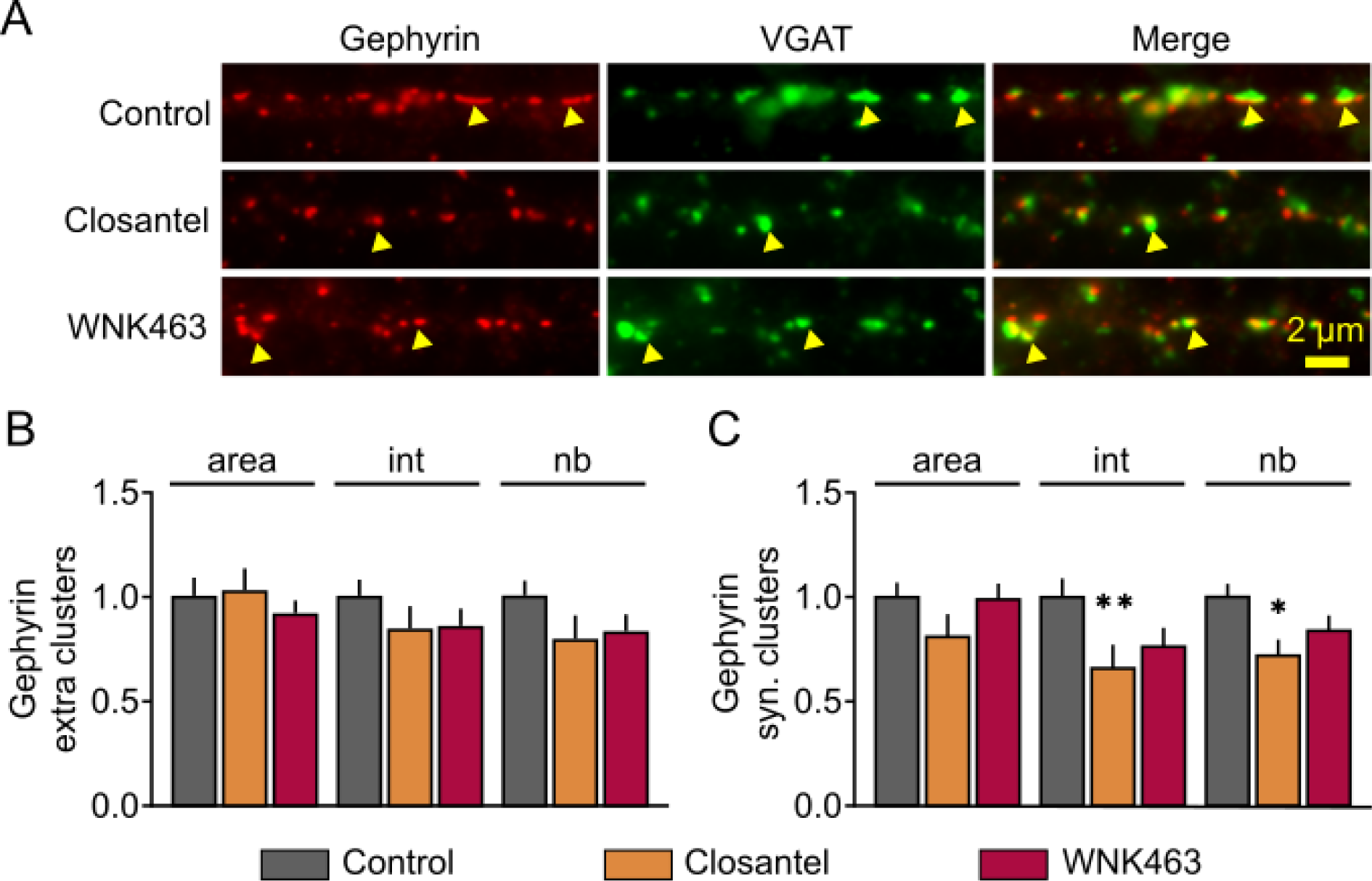
Blockade of the WNK pathway destabilizes synaptic gephyrin. **A.** Representative images of gephyrin (red), VGAT (green) and the merge (right) for control, closantel, and WNK463 treated neurons. Arrowheads show examples of synaptic clusters labeled for gephyrin and VGAT. Scale bar, 2 µm. **B-C.** Quantification of the mean surface area (area), intensity (int) and number (nb) of gephyrin clusters for extrasynaptic (B) and synaptic (C) clusters for control (grey), closantel (orange) and WNK463 (red) treatments. ***B:*** Control n=47 cells, Closantel n=31 cells, WNK463 n=48 cells. 4 cultures. Area: closantel p>0.9999, WNK463 p>0.9999, intensity: closantel p=0.4755, WNK463 p=0.7803, number: closantel p=0.1503, WNK463 p=0.7659; ***C:*** Control n=47 cells, Closantel n=31 cells, WNK463 n=48 cells. 4 cultures. Area: closantel p=0.1320, WNK463 p>0.9999, intensity: closantel p=0.0078, WNK463 p=0.3096, number: closantel p=0.0444, WNK463 p=0.1315. Data are presented as mean values ± SEM. Values were normalized to the corresponding control values. Kruskal-Wallis test.

### The WNK signaling pathway promotes gephyrin clustering by phosphorylating two key serine and threonine residues

Phosphorylation of gephyrin has been identified as a major regulatory mechanism for its clustering, as well as for the lateral diffusion and stabilization of GABA_A_Rs at inhibitory synapses (Tyagarajan et al., 2011, 2013; Battaglia et al., 2018). To identify residues potentially targeted by the WNK pathway, we performed a sequence analysis using a group-based phosphorylation prediction tool and selected candidate serine/threonine sites with probability scores above the medium confidence threshold. We identified eight candidate residues: S35 in the G domain; S232, T260, S280, and S303 in the central linker domain; and S325, S436, and S705 in the E domain **(Table 1).**

**Table 1.** Gephyrin residue candidates for WNK-mediated phosphorylation. Candidate residues are presented by C-ter to N-ter order. In bold, the candidate residue surrounded by the 7 upstream and downstream amino-acids. Only candidate residues with probability scores above the cut-off are shown. Related to Figure S3, Figure 4.

| Domain | Residue | Sequence | Probability Score | Cutoff |
| --- | --- | --- | --- | --- |
| G-domain | S35 | RNLAEDR**S**GINLKDL | 5.768 | 4.181 |
| C-domain (linker) | S232 | ASTEDSS**S**SHITAAA | 4.691 | 4.181 |
|  | T260 | VQVLPRDTASLSTTP | 5.351 | 4.181 |
|  | S280 | AQATSRL**S**TASCPTP | 4.69 | 4.181 |
|  | S303 | KENILRASHSAVDIT | 4.351 | 4.181 |
| E-domain | S325 | MSPFPLT**S**MDKAFIT | 5.009 | 4.181 |
|  | S436 | DTELIRE**S**DDGTEEL | 4.824 | 4.181 |
|  | S705 | SRLMSMR**S**ANGLLML | 5.211 | 4.181 |

To assess their functional relevance, we generated phosphomimetic gephyrin mutants by substituting these residues with glutamate and analyzed their effects on gephyrin clustering. Under basal conditions, overexpression of phosphomimetic mutants in any of the domains (G, linker, or E) did not significantly alter clustering compared to WT-gephyrin, as assessed by conventional fluorescence imaging **(Figure S3A-B).**

We next evaluated whether these constructs were sensitive to WNK pathway activation. As expected, activating WNK signaling by lowering intracellular chloride significantly increased WT-gephyrin cluster size and intensity, without changing cluster density **(Figure S3C).** This result aligns with the increased GABA_A_R clustering seen in low-chloride conditions **(Figure 2),** suggesting that **WNK activation promotes gephyrin-mediated recruitment of GABA_A_Rs to synapses.**

We then tested whether phosphomimetic mutants in the G (S35E) and E domains (S325E/S436E/S705E) responded to WNK activation. These mutants still exhibited increased clustering upon WNK activation **(Figure S3D-E),** indicating that these sites are not essential for WNK-mediated regulation of gephyrin. In contrast, a mutant combining several phosphosites in the central linker domain (S232E, T260E, S280E, S303E) failed to respond to WNK activation **(Figure S3F),** suggesting that one or more of these residues are functionally involved.

To refine this, we analyzed individual T260E and S280E mutants. Under basal conditions, neither mutant increased gephyrin clustering **(Figure S3G**). However, while WNK activation enhanced WT-gephyrin clustering **(Figure S3H),** it had no effect on T260E or S280E mutants **(Figure S3I-J),** implicating both residues in WNK-dependent regulation.

To test whether both residues are necessary and sufficient for this regulation, we generated a T260E/S280E double mutant. Although neither single mutant increased gephyrin clustering under basal conditions **(Figure S3G),** the double mutant significantly enhanced gephyrin clustering **(Figure 4A-B),** mimicking the effect of WNK activation on WT-gephyrin **(Figure 4C).** Furthermore, WNK activation no longer increased clustering of the double mutant **(Figure 4D),** confirming that phosphorylation of T260 and S280 is required for the effect.

**Figure 4.**
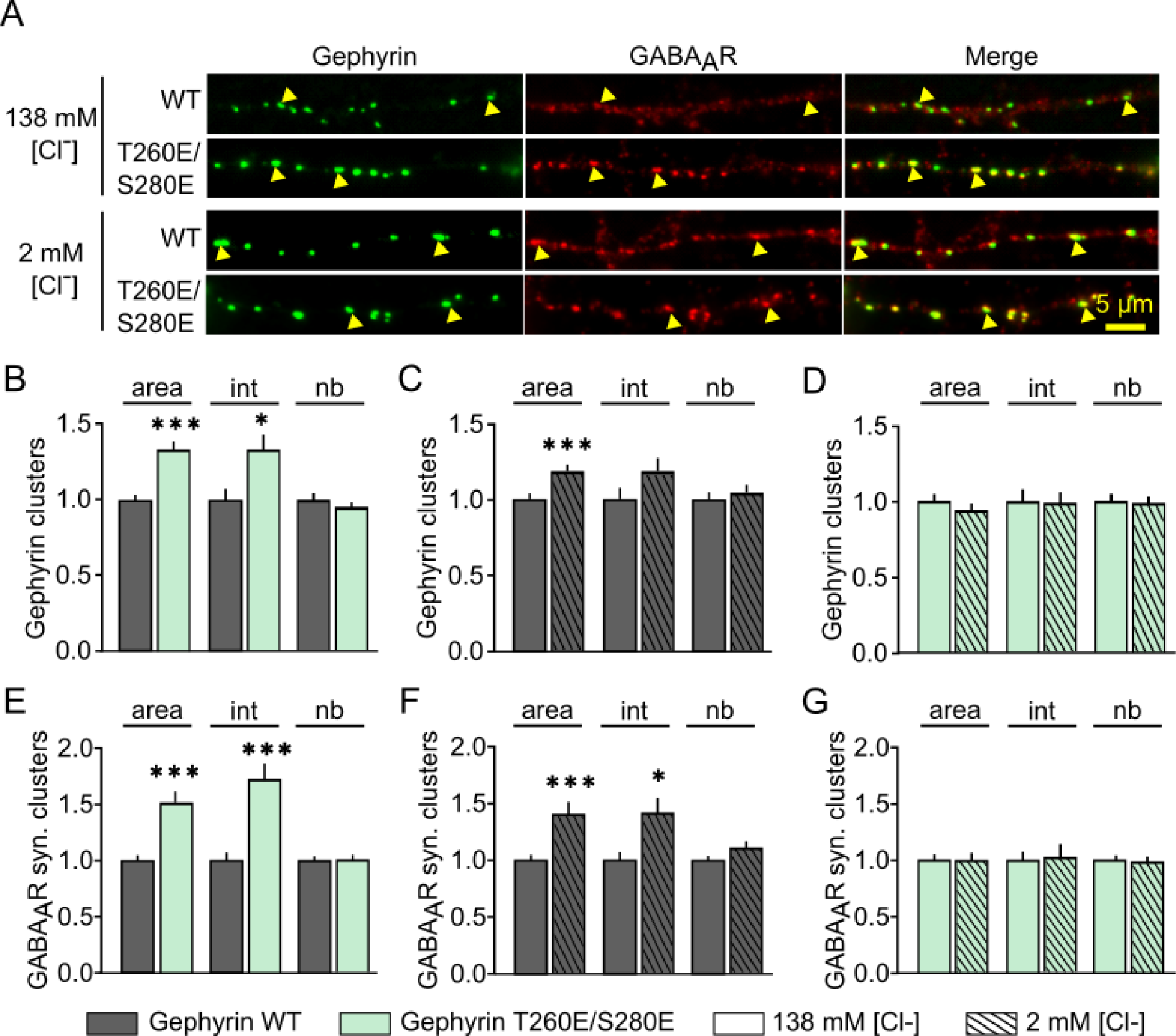
Phosphorylation of the Threonine 260 and Serine 280 residues of gephyrin increases GABA_A_R α1 accumulation in GABAergic synapses. **A.** Representative images of gephyrin (green), GABA_A_R α1 (red) and the merge (right) for neurons transfected with WT or T260E/S280E gephyrin and treated with 138 mM [Cl-] or 2mM [Cl-] solutions. Arrowheads show examples of clusters labeled for gephyrin and GABA_A_R α1. Scale bar, 5 µm. **B-G.** Quantification of the mean surface area (area), intensity (int) and number (nb) of gephyrin (B-D) and GABA_A_R α1 synaptic (E-G) clusters in neurons expressing gephyrin-WT (grey) or gephyrin-T260E/S280E (green) and treated with 138 mM [Cl-] (plain) or 2mM [Cl-] (hatched) solutions. WT n=71 cells, T260E/S280E n=75 cells, 138 mM [Cl-] n=65-71 cells, 2 mM [Cl-] n=60-64 cells, 7 cultures. ***B :*** area p<0.0001, Int p=0.0326, Nb p=0.6090; ***C :*** area p=0.0005, Int p=0.0719, Nb p=0.3770. ***D :*** area p=0.3324, Int p=0.8787, Nb p=0.3089; ***E :*** area p<0.0001, Int p=0.0002, Nb p=0.6689; ***F :*** area p=0.0001, Int p=0.0052, Nb p=0.6036; ***G :*** area p=0.2292, Int p=0.0739, Nb p=0.5214. Data are presented as mean values ± SEM. Values were normalized to the corresponding control values. Mann-Whitney test.

The double mutant had a similar effect on GABA_A_R α1: it increased receptor clustering under basal conditions **(Figure 4E),** comparable to WT after WNK activation **(Figure 4F),** and prevented additional recruitment of surface GABA_A_R α1 following WNK activation **(Figure 4G). Altogether, these data demonstrate that phosphorylation of gephyrin on T260 and S280 is both necessary and sufficient for WNK-mediated stabilization of gephyrin and GABA_A_R α1 at GABAergic synapses.**

### Functional impact of the WNK pathway *in vitro*

Reduced lateral diffusion and increased clustering of GABA_A_Rs at inhibitory synapses are known to enhance GABAergic transmission (Bannai et al., 2009; de Luca et al., 2017; Petrini et al., 2014; Petrini and Barberis, 2014). To evaluate the functional consequences of WNK-mediated stabilization of GABAergic synapses, we first conducted calcium imaging experiments in cultured hippocampal neurons. These experiments revealed that the WNK pathway modulates neuronal activity: acute inhibition of WNK signaling rapidly (within minutes) increased intracellular calcium levels ([Ca²⁺]_i_; **Figure S4A-B**), whereas pathway activation under low-chloride conditions decreased [Ca²⁺]_i_ **(Figure S4C-D).** This reduction in neuronal activity upon WNK activation is consistent with enhanced GABA_A_R diffusion-trapping and synaptic accumulation, supporting the hypothesis that WNK pathway activation strengthens inhibitory GABAergic signaling. Furthermore, activation of GABA_A_Rs with muscimol reduced [Ca²⁺]_i_ in both gephyrin-WT and T260E/S280E-expressing neurons **(Figure S4E-F).** After 15 minutes of treatment, the reduction was more pronounced in neurons expressing the T260E/S280E double mutant compared to those expressing WT gephyrin (WT: ΔF/F₀ = 0.71 ± 0.10; T260E/S280E: ΔF/F₀ = 0.46 ± 0.09; t-test, p = 0.0827). These results suggest that **neurons expressing the phosphomimetic gephyrin mutant may experience stronger GABAergic inhibition.**

### Functional and behavioral impact of the T260E/S280E gephyrin mutant *in vivo*

We next examined the *in vivo* effects of the gephyrin T260E/S280E double mutant. To this end, we performed bilateral injections of AAVs encoding either gephyrin-GFP-T260E/S280E or gephyrin-GFP-WT under the CamkII promoter into the hippocampus of adult mice **(Figure 5A).** Expression levels of gephyrin, GABA_A_R α1, and GABA_A_R α2 in the CA1 region were then assessed by immunohistochemistry and confocal microscopy **(Figure 5B).** Neurons expressing the T260E/S280E mutant showed increased GABA_A_R α1 immunoreactivity and a trend towards reduced GABA_A_R α2 immunoreactivity **(Figure 5C–E).**

**Figure 5.**
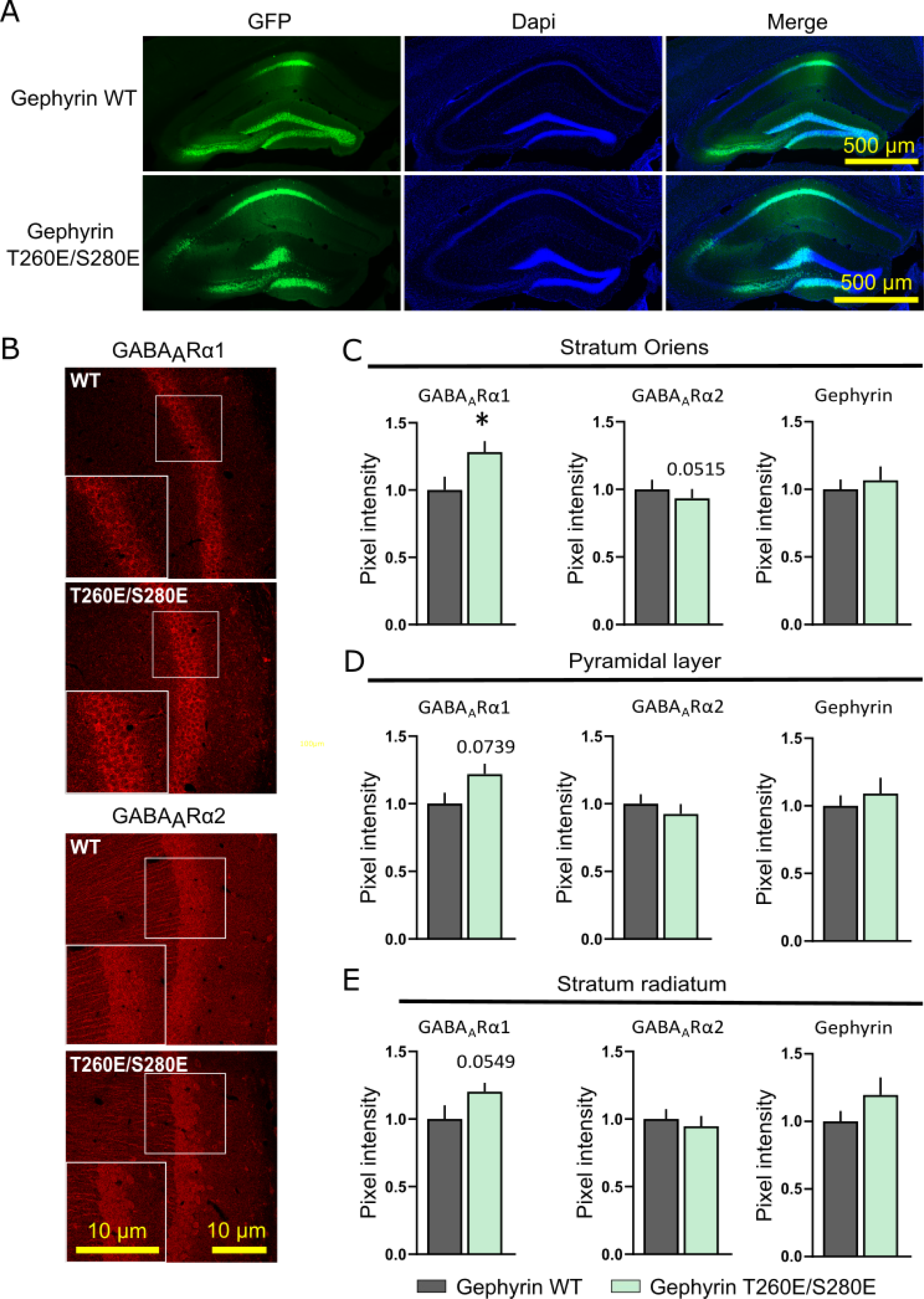
Expression of gephyrin-T260E/S280E increases GABA_A_R α1 immunoreactivity in the CA1 hippocampus. **A.** Representative hippocampal coronal sections immunostained for GFP (green) and DAPI (blue) from mice infected with AAVs of pCamKII-Gephyrin WT or pCamKII-Gephyrin T260E/S280E. Scale bar, 500 µm. **B**. Representative hippocampal coronal sections immunostained for GABA_A_R α1 or GABA_A_R α2 (red) in CA1 region from mice injected with AAVs of pCamKII-Gephyrin WT or pCamKII-Gephyrin T260E/S280E. Scale bar, 100 µm. **C-E.** Quantification of GABA_A_R α1, GABA_A_R α2 and gephyrin mean fluorescence intensity in region of interest drawn in the stratum oriens (C), pyramidal layer (D) and stratum radiatum (E) in Gephyrin WT (grey) and Gephyrin T260E/S280E (green) infected mice. WT n=10 animals, T260E/S280E n=13 animals. ***C:*** GABA_A_R α1 p=0.0461, GABA_A_R α2 p=0.0515, gephyrin p=0.5999; ***D:*** GABA_A_R α1 p=0.0739, GABA_A_R α2 p=0.4695, gephyrin. p=0.5294; **E:** GABA_A_R α1 p=0.0549, GABA_A_R α2 p=0.6193, gephyrin p=0.29004. Data are presented as mean values ± SEM. Values were normalized to WT group. Mann-Whitney or *t*-test. * *p* < 0.05.

Given that the subunit composition of GABA_A_Rs influences their deactivation kinetics and that α1-containing receptors deactivate faster than those containing α2 or α4 subunits (Eyre et al., 2012), we investigated whether the T260E/S280E mutation affected GABAergic currents. Consistent with a shift toward α1-containing receptors, miniature inhibitory postsynaptic currents (mIPSCs) decayed significantly faster in neurons expressing the T260E/S280E mutant compared to those expressing WT gephyrin **(Figure S5A–D).** This faster decay likely reflects increased synaptic incorporation of α1-containing GABA_A_Rs. These results, in agreement with our *in vitro* findings, support **a role for the WNK pathway in regulating the efficacy of GABAergic synapses *in vivo***.

To investigate whether *in vivo* expression of T260E/S280E-gephyrin in the hippocampus alters mouse behavior, we conducted a series of behavioral tests 10 to 14 days after AAV injection of either the mutant or WT construct. We first assessed general activity and social behaviors. Spontaneous locomotion in the actimeter showed no difference between T260E/S280E- and WT-expressing mice **(Figure S6A-B),** and no changes in stereotypic behaviors were observed in either the actimeter or marble burying tests **(Figure S6C-D).** Social interaction, measured by preference for an unfamiliar mouse over a neutral object, was also unaffected by mutant gephyrin expression **(Figure S6E-F).** In cognitive tasks, both groups performed similarly in place and object recognition tests, indicating intact spatial and object memory **(Figure S6G-J).** These findings suggest that **hippocampal expression of the T260E/S280E mutant does not impair locomotion, social behaviors or cognition.**

Given the established role of hippocampal GABAergic transmission in regulating anxiety (Rudolph et al., 1999; Löw et al., 2000; Engin, 2023), we next assessed anxiety-like behaviors. In the open field (OF) test under anxiogenic lighting (∼300 lux), mice expressing T260E/S280E-gephyrin displayed increased overall locomotion compared to WT controls **(Figure 6A-B).** They also traveled farther and spent more time in the center of the arena **(Figure 6C-D),** although these effects were moderate relative to the increase in general activity **(Figure 6E),** suggesting a possible reduction in anxiety. To further evaluate this effect, we used a second anxiety paradigm—the elevated O-maze. Mice expressing the T260E/S280E mutant spent more time and covered greater distances in the open (anxiogenic) arms compared to controls **(Figure 6F-G),** reinforcing the observation of reduced anxiety-like behavior.

**Figure 6.**
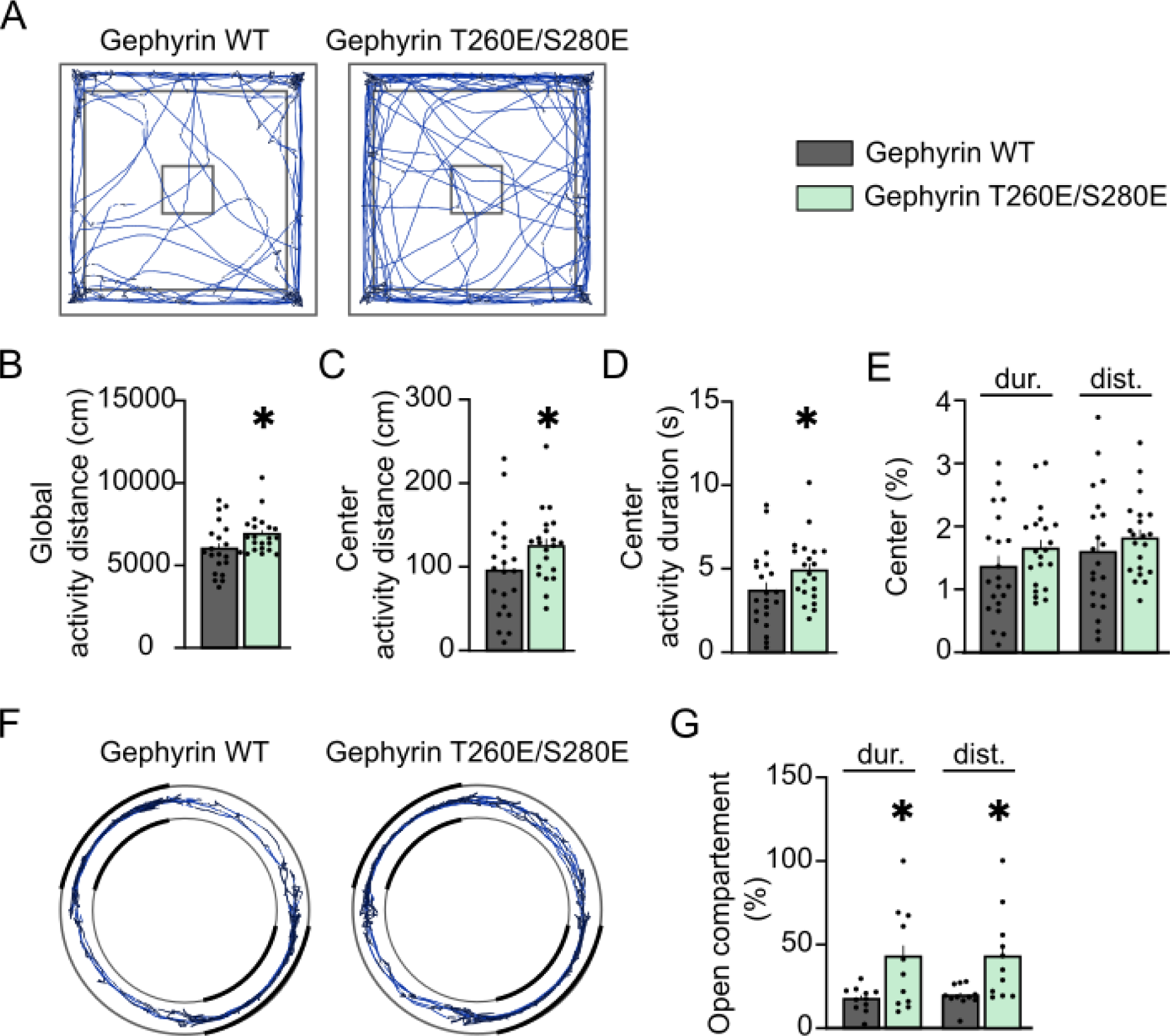
Hippocampal expression of Gephyrin-T260E/S280E decreases anxiety-like behavior in adult mice. **A.** Representative open field (OF) tracks (blue) for mice infected with pCamKII-Gephyrin WT (grey) vs. pCamKII-Gephyrin T260E/S280E(green) AAVs. **B-C.** Quantification of the distance of active exploration measured in the whole OF (B) or in the center of the OF (C) for gephyrin WT (grey) and T260E/S280E (green) infected mice. WT n=22 animals, T260E/S280E n=21 animals. ***B:*** p=0.0352; ***C:*** p=0.0398 **D.** Quantification of active exploration duration in the center of the OF for gephyrin WT and T260E/S280E infected mice. p=0.0398. **E.** Percentage of active exploration duration and distance in the center over active exploration in the whole OF for gephyrin WT (grey) and gephyrin T260E/S280E infected mice (light green). Duration p=0.2096, distance p=0.2501. **F.** Representative tracks (blue) in Elevated O-Maze (EOM) for mice infected with pCamKII-Gephyrin WT or pCamKII-Gephyrin T260E/280E AAVs. Light-gray zones represent open arms. **G.** Percentage of duration and distance in open compartment over exploration in the whole EOM for gephyrin WT and gephyrin T260E/S280E mice. WT n=11 animals, T260E/S280E n=11 animals. Duration p=0.0213, distance p=0.0109. Data are presented as mean values ± SEM. t-tests (B, C, E, G, I, J), Mann-Whitney (D).

**In summary, overexpression of T260E/S280E-gephyrin in the hippocampus decreases anxiety-like behavior in adult mice, without impairing general activity, social interaction or cognition. These results highlight a functional link between gephyrin phosphorylation, GABAergic synapse remodeling, and emotional behavior.**

## Discussion

In this study, we provide evidence for a regulatory mechanism of GABAergic synapse function in central hippocampal neurons that involves the chloride-sensitive WNK signaling pathway. Specifically, we show that WNK signaling regulates the diffusion–capture of GABA_A_Rs both at and outside synapses, as well as their membrane clustering, stability, and internalization. Blocking the WNK pathway promotes receptor endocytosis, whereas its activation stabilizes GABA_A_Rs at inhibitory synapses. At the synapse, this WNK-dependent regulation of GABA_A_R diffusion and clustering operates through modulation of the gephyrin scaffold *via* phosphorylation at two key serine and threonine residues in the central linker region of gephyrin, residues for which no function had been previously described. We further showed that manipulating WNK activity alters neuronal activity and modifies both the subunit composition and functional properties of synaptic GABA_A_Rs. Finally, our *in vivo* data reveal that phosphorylation of gephyrin at WNK-targeted sites has an anxiolytic effect, highlighting the physiological relevance of this regulatory mechanism.

### WNK signaling is active in mature neurons

WNK signaling plays a critical role during neuronal development by maintaining depolarizing GABAergic responses in the immature brain. This is achieved through the regulation of membrane stability of the cation-chloride cotransporters KCC2 and NKCC1, which are essential for establishing the chloride gradient and thereby determining GABA_A_R efficacy and polarity (Alessi et al., 2014; Friedel et al., 2015). While WNK activity declines as the central nervous system matures (Berglund et al., 2006; Friedel et al., 2015), it can be reactivated in pathological conditions characterized by impaired inhibition, such as epilepsy (Yang et al., 2013; Heubl et al., 2017; Jeong et al., 2018), ischemia (Bhuiyan et al., 2017; Zhang et al., 2020) and schizophrenia (Fernandez-Enright et al., 2014). However, work from our laboratory has shown that the WNK pathway remains active in mature hippocampal neurons both *in vitro* and *in vivo*, where it regulates the membrane availability, diffusion, clustering, and internalization of KCC2 and NKCC1, thereby influencing their function (Côme et al., 2023; Heubl et al., 2017; Pol et al., 2023). Our current study extends these findings by demonstrating that WNK signaling also directly modulates GABA_A_Rs and their scaffold protein gephyrin in mature hippocampal neurons, ultimately shaping GABAergic inhibitory transmission. Notably, the observation that pathway activation exerts a stronger effect on GABA_A_R lateral diffusion at synapses than pathway inhibition suggests that WNK signaling may be dynamically recruited under specific physiological or pathological conditions.

### WNK-dependent phosphorylation of gephyrin residues T260 and S280

Gephyrin phosphorylation plays a central role in organizing and compacting the subsynaptic gephyrin scaffold, which in turn governs the clustering of both gephyrin and GABA_A_Rs at inhibitory synapses (Tyagarajan et al., 2011, 2013; Kalbouneh et al., 2014; Flores et al., 2015; Battaglia et al., 2018). Given the destabilizing effect of WNK/SPAK inactivation on gephyrin clustering, we hypothesized that gephyrin may be a direct substrate of WNK/SPAK kinases. Bioinformatic analysis revealed potential WNK kinase phosphorylation sites within the main isoform of gephyrin. Supporting this prediction, our data show that phosphomimetic gephyrin (T260E/S280E) reproduces the WNK-mediated stabilization of GABAergic synapses. Notably, the serine/threonine kinases WNK, SPAK, and OSR1 are known to target multiple nearby phosphorylation sites on proteins such as NKCC1 (T203/T207/T212) and KCC2 (T906/T1007) (Inoue et al., 2012; Thastrup et al., 2012; de los Heros et al., 2014; Friedel et al., 2015). This makes the dual phosphorylation of gephyrin at T260 and S280 highly plausible, particularly given that these two residues are only 18 amino acids apart and reside within a flexible linker domain known to accommodate rapid conformational changes (Sander et al., 2013). However, it remains unclear whether both sites are phosphorylated simultaneously during a single kinase binding event, or whether phosphorylation of one site facilitates access to the other *via* conformational changes in the gephyrin molecule. Further structural and phosphoproteomic analyses are needed to resolve this mechanism in detail. While our findings demonstrate that phosphorylation at T260 and S280 is dependent on WNK signaling, it is possible that additional kinases downstream of WNK contribute to this regulation. The WNK pathway is known to interact with other signaling cascades involved in GABAergic synapse modulation, including the PI3K/Akt/mTOR pathway (Vitari et al., 2004; Jiang et al., 2005; Nishida et al., 2012; Yoshizaki et al., 2015; Saha et al., 2022; Sengupta et al., 2013) and the GSK3β pathway (Sato and Shibuya, 2018; Shimizu and Shibuya, 2022). However, we were able to exclude the involvement of GSK3β in our observations, as a gephyrin phosphorylation mutant specific to this pathway (S270A) still responded to WNK modulation (data not shown). Moreover, gephyrin integrates signals from multiple pathways *via* phosphorylation at distinct residues, which can act synergistically to fine-tune synaptic organization and function (Tyagarajan et al., 2013, 2011). Thus, a more comprehensive mechanistic analysis is necessary to delineate the full spectrum of gephyrin regulation by WNK signaling and its interactions with other pathways. Importantly, this study provides the first evidence of direct regulation of the GABAergic synapse by the WNK/SPAK signaling pathway.

### WNK signaling shapes GABAergic synaptic function and modulates behavior

We demonstrated that activation of the WNK signaling pathway, *via* phosphorylation of gephyrin, promotes the recruitment of GABA_A_R α1 subunits to inhibitory synapses. As increased synaptic receptor density generally enhances GABAergic transmission, this recruitment likely contributes to stronger inhibition. Consistent with this, calcium imaging in cultured neurons revealed that pharmacological activation of the WNK pathway reduced neuronal activity, while its inhibition increased it. Furthermore, neurons expressing the phosphomimetic T260E/S280E-gephyrin construct exhibited a greater response to muscimol compared to those expressing WT gephyrin, suggesting enhanced GABAergic inhibition. These findings indicate that the observed effects of WNK pathway modulation on neuronal activity primarily result from direct regulation of gephyrin and GABA_A_R stabilization at inhibitory synapses, rather than from off-target or indirect effects of the pharmacological agents used.

To further investigate the functional consequences of this mechanism, we used AAV-mediated expression of either WT or T260E/S280E-gephyrin *in vivo*. Expression of the phosphomimetic mutant significantly altered the kinetics of miniature inhibitory postsynaptic currents (mIPSCs), reducing the decay time constant. This acceleration in mIPSC decay likely reflects changes in receptor subunit composition, favoring faster-desensitizing α1-containing GABA_A_Rs (Bartos et al., 2001; Eyre et al., 2012). These electrophysiological data are in line with our *in vitro* and *in vivo* results showing WNK/SPAK-dependent recruitment of α1 subunits to synapses.

Together, these findings support the idea that WNK signaling plays a homeostatic role in the regulation of inhibitory synaptic strength. We propose that when neuronal chloride levels drop—such as after GABA_A_R closure or desensitization—the chloride-sensitive WNK pathway becomes activated. This activation could restore inhibitory tone by enhancing gephyrin-mediated clustering of GABA_A_Rs, thereby maintaining synaptic efficacy. Such a mechanism would act in parallel with WNK-mediated regulation of chloride transporters: WNK activation promotes the stability and membrane accumulation of the chloride importer NKCC1 (Côme et al., 2023), while downregulating the chloride exporter KCC2 (Heubl et al., 2017), thus elevating intracellular chloride. Overall, our work identifies the WNK pathway as a central regulator of the inhibitory synapse, orchestrating both chloride homeostasis and postsynaptic receptor composition and organization *via* gephyrin.

Several neurological disorders characterized by impaired inhibitory transmission in mature neurons are associated with elevated WNK pathway activity. As a result, therapeutic approaches have primarily aimed at dampening this pathway using pharmacological inhibitors of WNK/SPAK/OSR1, yielding promising outcomes in models of epilepsy (Yang et al., 2013; Jeong et al., 2018; Lee et al., 2022), ischemia (Bhuiyan et al., 2017; Zhang et al., 2020), and schizophrenia (Fernandez-Enright et al., 2014). In contrast, our study—alongside recent findings by Jaykumar et al. (2024)—points to a potentially beneficial role of WNK pathway activation in the adult hippocampus under physiological conditions. Notably, the recent report that WNK463 induces anxiogenic effects (Jaykumar et al., 2024) aligns with our own data showing that the phosphomimetic gephyrin mutant T260E/S280E exerts an anxiolytic effect. Together, these results suggest that controlled activation of the WNK pathway may offer a novel therapeutic strategy for treating anxiety disorders.

## Methods

### Experimental model and subject details

#### Animals

For all experiments performed on animals, procedures were carried out according to the European Community Council directive of 24 November 1986 (86/609/EEC), the guidelines of the French Ministry of Agriculture and the Direction Departmental de la Protection des Populations de Paris (Institut du Fer à Moulin, Animalerie des Rongeurs, license C 72-05-22). All efforts were made to minimize animal suffering and to reduce the number of animals used. Timed pregnant Sprague-Dawley rats were supplied by Janvier Lab (Le Genest St. Isle, France) and embryos were used at embryonic day 18 or 19 as described below. C57BL/6JRj mice, supplied by Janvier Lab, were delivered to our animal facility at least a week before surgery. Animals were housed in standard laboratory cages on a 12-hours light/dark cycle, in a temperature-controlled room (21 °C) with free access to food and water.

#### Dissociated hippocampal cultures

Primary cultures of hippocampal neurons were prepared as previously described (Chamma et al., 2013). Briefly, hippocampi were dissected from embryonic day 18 or 19 Sprague-Dawley rats of either sex. Tissue was then trypsinized (0.25% v/v), and mechanically dissociated in 1X HBSS (Invitrogen, Cergy Pontoise, France) containing 10 mM HEPES (Invitrogen,). Neurons were plated at a density of 180 × 10^3^ cells/mL onto 18-mm diameter glass coverslips (Assistent, Winigor, Germany) pre-coated with 50 μg/mL poly-D,L-ornithine (Sigma-Aldrich, St. Louis, USA) in plating medium composed of Minimum Essential Medium (MEM, Invitrogen) supplemented with horse serum (10% v/v, Invitrogen), L-glutamine (2 mM) and Na^+^ pyruvate (1 mM) (Invitrogen). After attachment for 3–4 h, cells were incubated in culture medium that consists of Neurobasal medium (Invitrogen) supplemented with B27 (1X) (Invitrogen), L-glutamine (2 mM) (Invitrogen), and antibiotics (penicillin 200 units/mL, streptomycin, 200 μg/mL) (Invitrogen) for up to 4 weeks at 37°C in a 5% CO_2_ humidified incubator. Each week, one-third of the culture medium volume was renewed.

### Method details

#### DNA constructs

The following constructs were used: pCAG_GPHN.FingR-eGFP-CCR5TC (Gross et al., 2013) was a gift from Don Arnold (Addgene plasmid # 46296; RRID:Addgene_46296), Gphn 3’-UTR shRNA (Yu et al., 2007) and EGFP-gephyrin P1 variant (Lardi-Studler et al., 2007). pRP-eGFP:RatGphn(S35E) (VB220621-1389pqf), pRP-eGFP:RatGphn (S325E, S436E, S705E) (VB220621-1403njc), pRP-eGFP:RatGphn (S232E, T260E, S280E, S303E) (VB220621-1404hwt), pRP-eGFP:RatGphn (T260E,S280E) (VB220310-1039vsu), pAAV-pCaMKII-eGFP:rGphn:WPRE (VB230323-1701pse) and pAAV-pCaMKII-eGFP:rGphn(T260E,S280):WPRE (VB230323-1723nmb) constructs were generated by Vector Builder (Chicago, USA) using the eGFP-gephryin P1 variant as template for site-directed mutagenesis.

#### Neuronal transfection

Transfections were carried out at DIV 13–14 using Transfectin (BioRad, Hercules, USA), according to the manufacturers’ instructions (DNA:transfectin ratio 1 μg:3 μL), with 0.5–1 μg of plasmid DNA per 20 mm well. Simple transfections of Gephyrin-FingR-eGFP were carried out with a plasmid concentration of 0.5 µg for SPT experiments. For cluster colocalization experiments, gephyrin mutants were always co-transfected with Gphn 3’-UTR shRNA with a 0.5:0.5 µg ratio. Experiments were performed 7 to 9 days post-transfection, on mature hippocampal neurons.

#### Pharmacology

All experiments were carried on DIV21-23 hippocampal neurons in conditions of neurotransmitter release and glutamatergic activity blockade by applying acutely the metabotropic glutamate receptor antagonist (S)-α-methyl-4-carboxyphenyl-glycine (MCPG; 500 μM; HelloBio, Bristol, UK), the ionotropic glutamate receptor antagonist 4-hydroxy-quinoline-2-carboxylic acid (kynurenic acid; 1 mM; Abcam, Cambridge, UK) and the voltage-gated sodium channels blocker tetrodotoxin (TTX; 1 μM; HelloBio).

The activation of the WNK pathway was favored by a 2 mM [Cl^−^] extracellular solution (1 mM CaCl_2_, 2 mM CH_3_SO_3_K, 2 mM MgSO_4_, 10 mM HEPES, 20 mM Glucose, 144 mM CH_3_SO_3_Na, pH 7,4). compared to a control 138 mM [Cl-] extracellular solution (2 mM CaCl_2_, 2 mM KCl, 3 mM MgCl_2_, 10 mM HEPES, 20 mM Glucose, 126 mM NaCl, 15 mM CH_3_SO_3_Na, pH 7,4). The blockade of the WNK pathway was favored by adding the pan-WNK inhibitor WNK463 (10 μM; MedChem Express, Monmouth Junction, USA) or the specific SPAK inhibitor, closantel (10 µM, Sigma, St. Louis, USA) either to the culture medium or to the 2 mM [Cl-] extracellular solution. In some experiments, the formation of endocytic zones was prevented by disrupting the interaction between dynamin and amphiphysin, an interaction that is essential for clathrin-coated pit-mediated endocytosis. For this purpose, cultured hippocampal neurons were pre-incubated with a 10 amino acid peptide (25 μM, Tocris, Bristol, UK) that blocks endocytosis together with drugs blocking the WNK pathway.

For single particle tracking experiments, neurons were transferred to a recording chamber and were pre-incubated for 10 min at 33°C with the drugs directly added to the imaging medium before starting the recordings. The imaging medium contained MEM without phenol red (Invitrogen,) limiting the auto-fluorescence and was supplemented with glucose (33 mM; Sigma), HEPES (20 mM) (Invitrogen), glutamine (2 mM) (Invitrogen), sodium pyruvate (1 mM) (Invitrogen) and B27 (1X) (Invitrogen). For calcium imaging, cells were loaded with Fluo4-AM (Life Technologies, Waltham, USA), transferred to a recording chamber, imaged in imaging medium first in absence of drugs for 300 s and then in presence of the appropriate drugs for another 600 s. For the immunocytochemistry, drugs were added to the culture medium 1 h prior fixation, in an incubator at 5% CO_2_ and at 37°C.

#### Live cell staining for single particle tracking

Neurons were stained as described previously (Merlaud et al., 2022). Briefly, cells were incubated for 10 min at 37°C with rabbit primary antibodies against extracellular epitopes of GABA_A_R α1 subunit (2 µg/ml, 224203, Synaptic Systems, Göttingen, Germany) and washed. Cells were then incubated for 1 min 37°C with anti-rabbit F(ab’)2-Quantum Dots (QD) emitting at 655 nm (1 nM; Invitrogen, 10592815) in PBS (1X; Invitrogen) in the presence of 10% Casein (v/v, Sigma) to prevent non-specific binding. Washing and incubation steps were done in imaging medium.

#### Single particle tracking and analysis

Cells were imaged using an Olympus IX71 inverted microscope equipped with a 60X objective (NA 1.42; Olympus, Tokyo, Japan) and an X-Cite 120Q (Excelitas Technologies Corp., Waltham, USA). Individual images of Gephyrin-FingR-eGFP and QD real time recordings (integration time of 75 ms over 1200 consecutive frames) were acquired with Hamamatsu ImagEM EMCCD camera (Hamamatsu, Japan) and MetaView software (Meta Imaging 7.7, Molecular Devices, USA). Cells were imaged within 50 min following labeling.

QD tracking and trajectory reconstruction were performed with homemade software (MATLAB; The Mathworks, Natick, USA) as described in Bannai et al., 2006. One sub-regions of dendrites were quantified per cell. In cases of QD crossing, the trajectories were discarded from analysis. Trajectories were considered synaptic when overlapping with the synaptic mask of gephyrin clusters or its perisynaptic zone of 2 pixels (380 nm). Out of this vicinity, trajectories were considered extrasynaptic. Values of the mean square displacement (MSD) plot versus time were calculated for each trajectory by applying the relation:

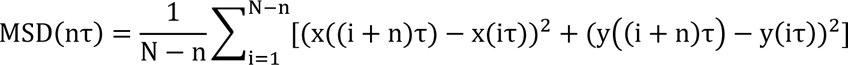

where τ is the acquisition time, N the total number of frames, n and i positive integers with n determining the time increment (Saxton and Jacobson, 1997). Diffusion coefficients (D) were calculated by fitting the first four points without origin of the MSD versus time curves with the equation: where b is a constant reflecting the spot localization accuracy. The explored area of each trajectory was defined as the MSD value of the trajectory at two different time intervals of at 0.42 and 0.45 s (Renner et al., 2012). Synaptic dwell time was defined as the duration of detection of QDs at synapses on a recording divided by the number of exits as detailed previously. Dwell times ≤5 frames were not retained.

Experimenters were blind to the condition of the sample analyzed. Sample size selection for experiments was based on published experiments, pilot studies as well as in-house expertise.

#### Immunocytochemistry

After being exposed for 1 h at 37°C to the appropriate drugs, cells were fixed for 5 min at room temperature (RT) in paraformaldehyde (PFA, 4% w/v, Sigma) and sucrose (14% w/v, Sigma) solution prepared in PBS (1X). We then stained GABA_A_R α1, gephyrin and or VGAT following different portocols, detailed below. Sets of neurons compared for quantification were labeled simultaneously.

##### Staining for surface/total experiments

Cells were incubated for 30 min minimum at RT in normal goat serum (GS) (10%, v/v, Invitrogen) in PBS to block non-specific staining. GABA_A_R α1 present at the neuronal surface were labeled by incubating cells for 1h at RT with primary antibodies against an extracellular epitope of the GABA_A_R α1 subunit (rabbit: 0,5 μg/mL, 224203, Synaptic Systems) prepared in PBS-GS 3% solution. After three washes in PBS (1X), cells were incubated for 45 min at RT with CY3-conjugated goat anti-rabbit antibodies (3.75 μg/mL, 111-095-003, Jackson Immunoresearch, West Grove, USA) prepared in PBS-GS 3% solution. After three washes in PBS (1X), cells were permeabilized for 4 min at RT with Triton X-100 (0.25% w/v; Invitrogen) in PBS (1X). Cells were subsequently incubated for 1 h at RT with rabbit GABA_A_R α1 primary antibodies (0,5 μg/mL, 224203, Synaptic Systems), in PBS-GS 3% solution to labelled total GABA_A_R α1. After three washes in PBS (1X), cells were incubated for 45 min at RT with Alexa 488 conjugated donkey anti-rabbit secondary antibodies (1.875 μg/mL, 715-547-003 Jackson Immunoresearch) to reveal total GABA_A_R α1. After three washes in PBS (1X), cells were finally mounted on glass slides using Mowiol 4–88 (48 mg/mL, Sigma).

##### Staining for STORM experiments

Cells were incubated for 30 min minimum at RT in normal GS (10%, v/v, Invitrogen). GABA_A_R α1 present at the neuronal surface were labeled by incubating cells for 1h at RT with primary antibodies against an extracellular epitope of the GABA_A_R α1 subunit (rabbit: 0,5 μg/mL, 224203, Synaptic Systems,) prepared in PBS-GS 3% solution. After three washes, cells were incubated for 45 min at RT with Alexa 647 conjugated goat anti-rabbit secondary antibodies (1.875 μg/mL, 111-605-003, Jackson Immunoresearch) were used to reveal surface GABA_A_R α1. After three washes in PBS (1X), cells were kept in PBS 1X until imaging.

##### Staining for colocalization experiments

For endogeneous gephyrin and VGAT colocalization, cells were permeabilized for 4 min at RT with Triton X-100 (0.25% w/v; Invitrogen) in PBS (1X). Cells were incubated with a mix of mouse gephyrin antibodies (5 μg/mL, 147011, Synaptic Systems,) and rabbit VGAT antibodies (1µg/mL, 131011, Synaptic Systems) in PBS-GS 3% solution. After three washes, cells were incubated for 45 min at RT with a mix of Alexa 488 conjugated donkey anti-mouse antibodies (1.875 μg/mL,715-545-150, Jackson Immunoresearch) and CY3-conjugated goat anti-rabbit secondary antibodies (3.75 μg/mL, 111-095-003, Jackson Immunoresearch). After three washes, cells were finally mounted on glass slides using Mowiol 4–88 (48 mg/mL, Sigma).

For endogenous GABA_A_R α1 and gephyrin-GFP colocalization, cells were incubated for 30 min minimum at RT in normal GS (3%, v/v, Invitrogen) in PBS. GABA_A_R α1 present at the neuronal surface were labeled by incubating cells for 1h at RT with primary antibodies against an extracellular epitope of the GABA_A_R α1 subunit (rabbit: 0,5 μg/mL, 224203, Synaptic Systems) prepared in PBS-GS 3% solution. After three washes, cells were incubated for 45 min at RT with CY3-conjugated goat anti-rabbit antibodies (3.75 μg/mL, 111-095-003, Jackson Immunoresearch, West Grove, USA) or prepared in PBS-GS 3% solution. After three washes, cells were permeabilized for 4 min at RT with Triton X-100 (0.25% w/v; Invitrogen) in PBS (1X). After three washes, cells were finally mounted on glass slides using Mowiol 4–88 (48 mg/mL, Sigma).

#### STORM acquisition and analysis

STORM imaging was carried out on an inverted N-STORM Nikon Eclipse Ti microscope (Tokyo, Japan) with a 100x oil-immersion objective (N.A. 1.49) and an Andor iXon Ultra 897 EMCCD camera (image pixel size, 160 nm) (Oxford Instruments, Abingdon, UK), using specific lasers for STORM imaging of Alexa 647 (640 nm). Samples were imaged in a PBS-based oxygen-scavenging buffer containing imaging buffer (Tris 100 mM Invitrogen, NaCl 20 mM, pH 8), glucose 40% (w/v, Sigma-Aldrich), PBS 1X, cysteamine hydrochloride (77 mg/mL, Sigma-Aldrich), catalase 5 mg/mL (Sigma-Aldrich) and glucose oxidase 67.6 U/mL (Sigma-Aldrich). Catalase was diluted in MgCl 4 mM, EGTA 2 mM, and PIPES 24 mM (Sigma-Aldrich, pH 6.8). Pyranose oxidase was diluted in the same buffer supplemented with glycerol (50% v/v). Movies of 30 000 frames were acquired at frame rates of 20 ms. A Nikon Perfect Focus System maintained the z position during the acquisition.

Single-molecule localization and 2D image reconstruction was conducted as described in Specht et al., 2013 by fitting the PSF of spatially separated fluorophores to a 2D Gaussian distribution. The drift of the stage was corrected using 100 nm multicolor fluorescent beads (TetraSpeck, 1:300, Invitrogen) to follow the movement through the frames; and was nullified by superimposing the molecular localizations of each frame. STORM images were rendered using Diinamic-R (Paupiah et al., 2023) by superimposing the coordinates of single-molecule detections, which were represented with 2D Gaussian curves of unitary intensity and SDs representing the localization accuracy (σ = 20 nm). The surface of GABA_A_R clusters and the densities of molecules per μm^2^ were measured in reconstructed 2D images through cluster segmentation based on detection densities. The minimum thresholds used to determine the clusters were: intensity=1%, detection density= 0.1 per nm^2^, number of detections=10, diameter for synaptic clusters: 150nm, diameter comprised between 50nm and 150nm for extrasynaptic clusters. The resulting binary image was analyzed with the function “regionprops” of Matlab to extract the surface area of each cluster identified using this function. The density was calculated as the total number of detections in the pixels belonging to a given cluster, divided by the area of the cluster.

Experimenters were blind to the condition of the sample analyzed. Sample size selection for experiments was based on published experiments, pilot studies as well as in-house expertise.

#### Fluorescence image acquisition and analysis

Image acquisition was performed using a 63 objective (NA 1.32) on a Leica (Nussloch, Germany) DM6000 upright epifluorescence microscope with a 12-bit cooled CCD camera (Micromax, Roper Scientific, Evry, France) run by MetaMorph software (Roper Scientific). Image exposure time was determined on bright cells to obtain best fluorescence to noise ratio and to avoid pixel saturation. All images from a given culture were then acquired with the same exposure time and acquisition parameters. For surface/total expression analysis, quantification was performed using ImageJ (National Institutes of Health and LOCI, University of Wisconsin, USA). Several dendritic regions of interest were manually chosen and the mean average intensity per pixel was measured. For cluster colocalization analysis, quantification was performed using the Meta Imaging 7.7 software (Molecular Devices, USA). For each image, several dendritic regions of interest were manually chosen. The images were then flattened, the background was filtered (kernel size, 3 × 3 × 2) to enhance cluster outlines, and a user-defined intensity threshold was applied to select clusters and avoid their coalescence. For quantification of GABA_A_R synaptic clusters, clusters comprising at least 2 pixels and colocalized on at least 1 pixel with gephyrin clusters were considered. For quantification of gephyrin synaptic clusters, clusters comprising at least 2 pixels and colocalized on at least 1 pixel with VGAT clusters were considered. The number of clusters, the surface area and the integrated fluorescence intensities of clusters were measured.

Experimenters were blind to the condition of the sample analyzed. Sample size selection for experiments was based on published experiments, pilot studies as well as in-house expertise.

#### Computational phosphorylation site prediction

The gephyrin sequence from the eGFP-gephryin P1 variant was screened for putative phosphorylation residue by WNK kinases using the Group-based Prediction system 2.1 program (Xue et al., 2011, 2008; Xue et al., 2005). Only prediction sites with a probability score superior to the GPS-defined medium threshold were tested.

#### Calcium imaging

Neurons at 21-24 DIV were loaded with 10 μM Fluo-4AM (14217, Life Technologies) for 5 min at 37°C in imaging medium. After washing excess dye, cells were further incubated for 5– 10 min to allow hydrolysis of the AM ester. All incubation steps and washes were performed in imaging medium.

Cells were imaged at 37°C in an open chamber mounted on an inverted N-STORM Nikon Eclipse Ti microscope (Tokyo, Japan) with a 100x or 63x oil-immersion objective (N.A. 1.49). Fluo4-AM was illuminated using 472 (± 30) nm light from a diode. Emitted light was collected using a 520 (± 35) nm emission filter. Time-lapse at 0.033 Hz were acquired with an exposure time of 80-100 ms for 15 min with an Andor iXon Ultra 897 EMCCD camera (image pixel size, 160 nm) (Oxford Instruments), using Nikon software. The analysis was performed on a section of the soma that was in focus at different time points. Fluorescence intensities collected in the soma before and following bath addition of the drugs, were background subtracted. The data were analyzed using ImageJ (NIH, USA). For each cell, the mean fluorescence intensity in each 5 min time bin was normalized by the respective mean fluorescence intensity before drug application.

Experimenters were blind to the condition of the sample analyzed. Sample size selection for experiments was based on published experiments, pilot studies as well as in-house expertise.

#### Stereotaxic surgery

Eight-week-old mice were anesthetized with isoflurane (4% for induction, 1–2% for maintenance) and placed on a heating pad at 36-37°C for the entire surgery. Thirty minutes before opening the skin, buprenorphine was administered (0.1 mg/kg s.c). Two to three minutes before opening the skin, lidocaine (4 mg/kg s.c) was applied beneath the scalp. The virus was injected bilaterally in the dorsal hippocampus (500 nL in both the dentate gyrus and the CA1 region for each hemisphere) at the following stereotaxic coordinates from Bregma: −1.8 mm anteroposterior (AP), ±1.2 mm mediolateral (ML) and −2.1 and −1.3 mm dorsoventral (DV). The anti-inflammatory metacam was administered right after surgery, and once a day for the next two days (10 mg/kg s.c). After surgery, the body temperature was maintained using a heating pad under the cage until the animal recovered from anesthesia. Behavioral experiments were started 10 to 14 days after surgery. The spread of viral infection was systematically verified at the end of the experiments. Only data from mice with generalized gephyrin-eGFP expression in dorsal CA1, CA3 and DG were considered for behavioral analysis.

#### *Ex-vivo* patch-clamp experiments

After behavioral tests, mice (14-22 weeks) were deeply sedated using ketamine/xylazine (150mg/kg and 10mg/kg i.p.). Mice were perfused transcardially with ice-cold oxygenated (95% O2 – 5% CO2) solution containing 110 mM choline chloride, 2.5 mM KCl, 25 mM glucose, 25 mM NaHCO3, 1.25 mM NaH2PO4, 0.5 mM CaCl2, 7 mM MgCl2, 11.6 mM L-ascorbic acid, and 3.1 mM sodium pyruvate. After fast extraction, brains were cut into 250 μm coronal slices in the same solution using a vibrating-blade microtome (HM 650 V, Microm International GmbH). Slices were then stored in an artificial cerebrospinal fluid (aCSF) solution containing 125 mM NaCl, 2.5 mM KCl, 25 mM glucose, 25 mM NaHCO3, 1.25 mM NaH2PO4, 2 mM CaCl2, and 1 mM MgCl2 continuously oxygenated (95% O2 – 5% CO2).

To perform *ex vivo* patch-clamp recordings, slices were transferred to the recording chamber and were continuously maintained at 30-32°C in oxygenated aCSF (95% O2 – 5% CO2). In the dentate gyrus, Gephyrin WT or Gephyrin T260E/S80E infected cells were recognized by their GFP expression on an upright microscope (Olympus BX5WI). Cells were recorded in whole-cell configuration and voltage-clamp mode using a borosilicate glass pipette (3-6 MΩ) (GC150TF-7.5, Warner Instruments) and a DMZ-Universal-Electrode-Puller (Zeitz). To record miniature inhibitory postsynaptic currents (mIPSCs), pipettes were filled with an intracellular solution containing: 60 mM Cs methanesulfonate, 70 mM CsCl, 10 mM Hepes, 10mM Phosphocreatine, 0.2 mM EGTA, 8 mM NaCl, 2 mM ATP-Mg (pH 7.25, adjusted with CsOH). mIPSCs were recorded at a holding membrane potential of −65mV in the presence of tetrodotoxin (TTX; 0.5μM, Hello Bio) to block action potential-dependent synaptic transmission as well as CNQX (10μM; Hello Bio) and D-AP5 (50μM; Hello Bio) to isolate inhibitory currents. Recordings were performed using an EPC-10 amplifier (HEKA Electronik). The liquid junction potential (−5mV) was left uncorrected. Signals were low pass-filtered at 4kHz and sampled at 20 kHz. mIPSCs recordings were filtered offline at 2 kHz and 100 s of each recording were analyzed using MiniAnalysis software (Synaptosoft). Experimenters were blind to the condition of the sample analyzed. Sample size selection for experiments was based on published experiments, pilot studies as well as in-house expertise.

#### Behavioral tests

Experimenters were blind to the condition of the groups analyzed. Group size selection for experiments was based on published experiments, pilot studies as well as in-house expertise. Four independent cohorts were used. The entire testing period spanned 3 to 4 weeks, with a minimum interval of 1 day between consecutive tests.

##### Locomotor activity

We used actimeter racks of 8 circular corridors each (Imetronic, Pessac, France) equipped with infrared sensors to detect and automatically record both horizontal (locomotion) and vertical (rearing) activities. Animals were introduced individually in corridors and their spontaneous novelty driven activity was recorded every 5 minutes for 2 consecutive hours.

##### Open field

The test took place in a white square open-field (1×1 m) apparatus under high illumination (< 300 lux). The central area of the open field was a square of 21 cm per side, while the peripheral zone bordering the walls was 5 cm wide. Animals were individually placed in a corner and were allowed to freely explore the open-field. A video tracking system, which included a computer-linked overhead camera, was used to monitor general locomotor activity and activity in the center every min for 9 consecutive minutes (ViewPoint, Lyon, France).

##### Elevated-O-maze

The test took place under high illumination (∼300 lux) in a white circular maze (diameter: 51.5 cm, lane width: 6 cm), located at 42 cm above the floor, and divided into 4 quadrants. Two opposite quadrants were surrounded by dark, opaque walls (closed arms) and two other opposite quadrants were without walls (open arms). The mice were placed in one of the closed arms and were considered to be in the open arms when all four paws were detected in the arm. A video tracking system, which included a computer-linked overhead camera, was used to monitor activity in the closed and open arms every min for 9 consecutive minutes (ViewPoint).

##### Marble burying

Mice were individually assessed in a Plexiglas cage (20 x 15 x 25 cm) filled with 3 cm sawdust, overlaid with 12 glass marbles arranged in 3 lines of 4 equidistant marbles. The number of visible marbles (> 25% visible) was scored by two observers every minute for the first 10 minutes of the test, and every 5 minutes for the following 20 minutes.

##### Social exploration

The test took place in a square arena (50 x 50 cm) apparatus under low illumination (< 50 lux). Mice were first allowed to habituate to the arena for 10 minutes, then habituated to the chamber containing the two empty small round wire pots for 10 minutes. For the final stage, an unfamiliar mouse was placed under one wire pot and an object under the other one. Mice were left to freely explore for 10 minutes. A video tracking system, which included a computer-linked overhead camera allowing nose detection, was used to monitor exploration of each wire pot every minute for each trial (ViewPoint, Lyon, France). A preference index (PI) was used as a measure of preference for the unfamiliar mouse as compared to the object and was obtained as follows:

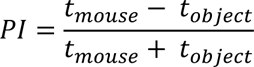

##### Place recognition and object recognition

Mice were tested successively for the detection of novel spatial position of an object and of a novel visual object. The test took place in a square arena (50 x 50 cm) apparatus under low illumination (< 50 lux). On the testing day, mice were placed in the arena and were allowed to explore three identical objects during 4 acquisition trials of 5 minutes each and separated by a 3 minutes ITI. On the fifth trial, 3 minutes later, mice were placed back into the arena in which one of the objects was moved to a new spatial position (place recognition). The mice were exposed again one more time to this new configuration of objects and on the seventh trial one object was substituted by a novel object with a different shape color and texture (object recognition). A video tracking system, which included a computer-linked overhead camera allowing nose detection, was used to monitor exploration of each wire pot every minute of each trial (ViewPoint, Lyon, France). A novelty index (NI) was used as a measure of discrimination between novel and old locations or novel and old objects and was obtained as follows:

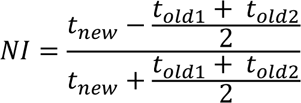

#### Histology

Mice were deeply sedated using ketamine/xylazine (150mg/kg and 10mg/kg i.p.) and then transcardially perfused with ice-cold 4% (w/v) paraformaldehyde (PFA) in PBS pH 7.4. Extracted brains were fixed in 4% PFA overnight and then equilibrated in 30% (w/v) sucrose in PBS. A sliding microtome HM450 Microm (Thermofisher, Walldorf, Germany) was used to section the brains at −35°C in 40 μm-thick coronal sections, collected in PBS and stored at −20°C in Amaral.

#### Immunohistochemistry

To verify the AAV injection sites, slices were preincubated for 15 minutes at room temperature in a solution containing 0.3% Triton X-100 (w/v; Invitrogen) in PBS (1X). Slices were then incubated for 1 hour minimum with slow agitation at RT in normal GS 10% (v/v, Invitrogen), 0.3% Triton X-100 (w/v; Invitrogen) in PBS (1X) to block non-specific staining. Incubation with chicken anti-GFP primary antibodies (2 µg/ml, AB16901, Millipore) was then performed in a 3% v/v (GS, Invitrogen), 0.3% Triton X-100 (w/v, Invitrogen) in PBS (1X) for 48 hours with slow agitation at 4 °C. After 3 x 10 min washes with 0.3% Triton X-100 (w/v; Invitrogen) in PBS (1X), slices were incubated with Alexa-488 conjugated donkey anti-chicken secondary antibodies (1.5 µg/ml, 703-545-155, Jackson Immunoresearch) in a 3% (v/v, GS, Invitrogen), 0.3% Triton X-100 (w/v, Invitrogen) in PBS (1X) for 5 hours with slow agitation at RT. After 3 x 10 min washes, slices were incubated in DAPI (1:2000, Sigma, MBD0015) in a 3% (v/v, GS, Invitrogen), 0.3% Triton X-100 (w/v, Invitrogen) in PBS (1X) for 20 minutes with slow agitation at RT. Finally, slices were washed 3 x 10 minutes and mounted in Mowiol 4–88 (48 mg/mL, Sigma).

Images of the whole hippocampus (Figure 5A) were acquired using a 5x objective on the automated microscope Cell Discoverer 7 (Zeiss, Oberkochen, Germany), equipped with an Axiocam 506m camera (Zeiss), run by Zen black software (Zeiss).

For *in vivo* quantification of gephyrin, GABA_A_R α1, or GABA_A_R α2 immunoreactivity, slices were incubated with sodium citrate 50mM pH 8.9 (an antigen retrieval) for 30 minutes at 80°C. Slices were cooled down at RT for 10 to 15 minutes. After 3 x 10 min washes with 0.3% Triton X-100 (w/v; Invitrogen) in PBS (1X), slices were incubated for 1 hour minimum with slow agitation at RT in normal GS 10% (v/v, Invitrogen), 0.3% Triton X-100 (w/v; Invitrogen) in PBS (1X) to block non-specific staining. Incubation with chicken anti-GFP primary antibodies (2 µg/ml, AB16901, Millipore), rabbit anti-GABA_A_R α1 (2 µg/ml, 224203, Synaptic Systems) or rabbit anti-GABA_A_R α2 (2 µg/ml, 224103, Synaptic Systems), mouse anti-gephyrin (2 µg/ml, 147011, Synaptic Systems) or mouse anti-VGAT (2 µg/ml, 131011, Synaptic Systems) primary antibodies was then performed in a 3% GS (v/v, Invitrogen), 0.3% Triton X-100 (w/v, Invitrogen) solution prepared in PBS (1X) for 48 hours with slow agitation at 4 °C. After 3 x 10 min washes with 0.3% Triton X-100 (w/v, Invitrogen) in PBS (1X), slices were incubated with Alexa-488 conjugated donkey anti-chicken (1.5 µg/ml, 703-545-155, Jackson Immunoresearch), CY3-conjugated goat anti-rabbit (1.5 μg/mL, 111-165-003, Jackson Immunoresearch), Alexa-647 conjugated donkey anti-mouse (1.5 μg/mL, 715-605-150, Jackson Immunoresearch), secondary antibodies in a 3% (v/v, GS, Invitrogen), 0.3% Triton X-100 (w/v, Invitrogen) solution prepared in PBS (1X) for 5 hours with slow agitation at RT. After 3 x 10 min washes, slices were incubated in DAPI (1:2000, Sigma, MBD0015) in a 3% (v/v, GS, Invitrogen), 0.3% Triton X-100 (w/v, Invitrogen) in PBS (1X) for 20 minutes with slow agitation at RT. Finally, slices were washed 3 x 10 minutes and mounted in Mowiol 4–88 (48 mg/mL, Sigma).

Image acquisition was performed using z-stack confocal imaging (0.5 µm step size) on either a Leica TCS SP5 upright confocal microscope or a Leica SP8 confocal microscope, both equipped with a 40x oil-immersion objective and operated with LAS X software (Leica). For quantification, three regions of interest (ROIs) were selected per hippocampal layer (stratum oriens, pyramidal layer, and stratum radiatum) in each of two coronal brain slices per animal. The mean gray intensity per pixel was measured using FIJI (ImageJ).

Experimenters were blind to the condition of the sample analyzed. Sample size selection for experiments was based on published experiments, pilot studies as well as in-house expertise.

### Statistics

For each experiment, quantifications are summed up in Table 2 and Table S1. Unless otherwise stated, normally distributed data are presented as mean ± SEM (standard error of the mean), whereas non-normally distributed data are given as medians ± IQR (inter quartile range). For each experiment, statistical details (n, p-values, and test used) can always be found in the figure legends. All statistical analyses were performed using GraphPad Prism 10.1.0 (Dotmatics, Boston, USA). Normality of distribution was assessed by the Shapiro-Wilk test. Normally distributed unpaired datasets were compared using the unpaired t test, ordinary one-way ANOVA tests followed by Holm-Sidak’s multiple comparison tests or two-way ANOVA tests followed by Sidak’s multiple comparison for kinetic analyses. Non-Gaussian unpaired datasets were tested by two-tailed unpaired non-parametric Mann-Whitney test or Kruskal– Wallis tests followed by Dunn’s multiple comparison tests. Normally distributed paired datasets were compared using paired t-test, whereas non-Gaussian paired datasets were tested by the paired Wilcoxon test. Cumulative distributions were compared with the Kolmogorov-Smirnov test. Indications of significance corresponding to p-values < 0.05 (*), p < 0.01 (**), p < 0.001 (***) are reported in the figures and in the corresponding legends.

**Table 2.**
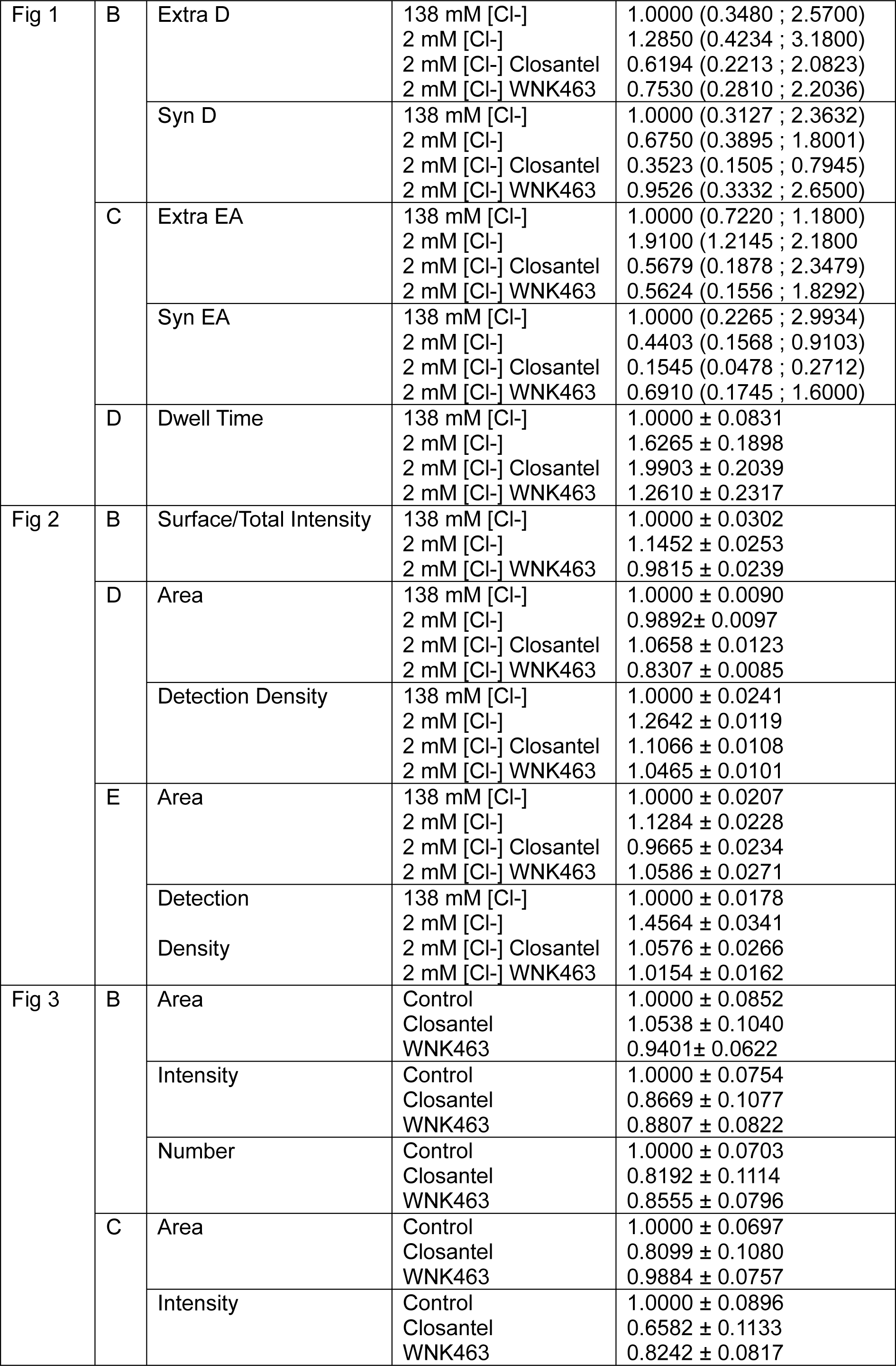
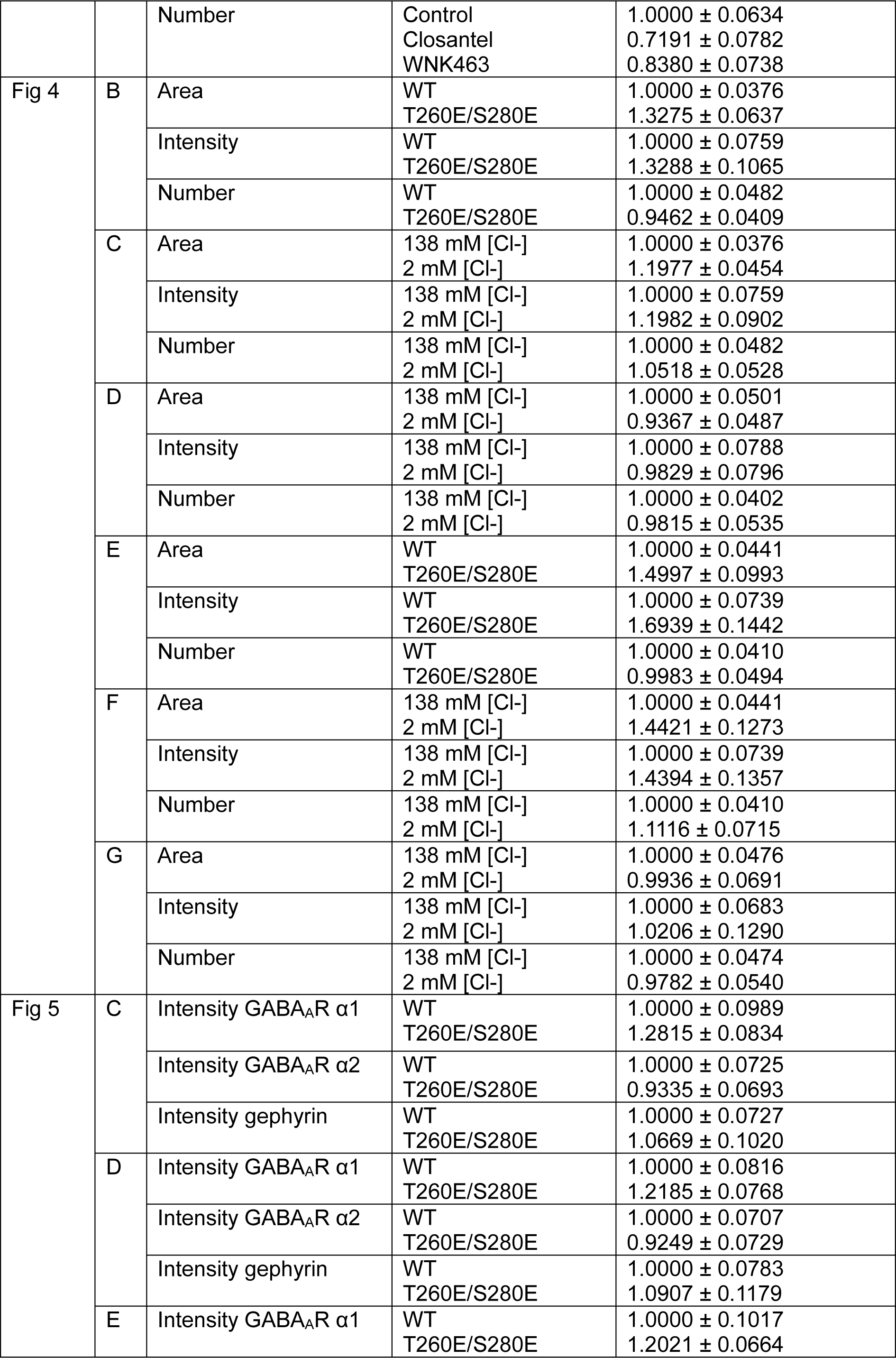
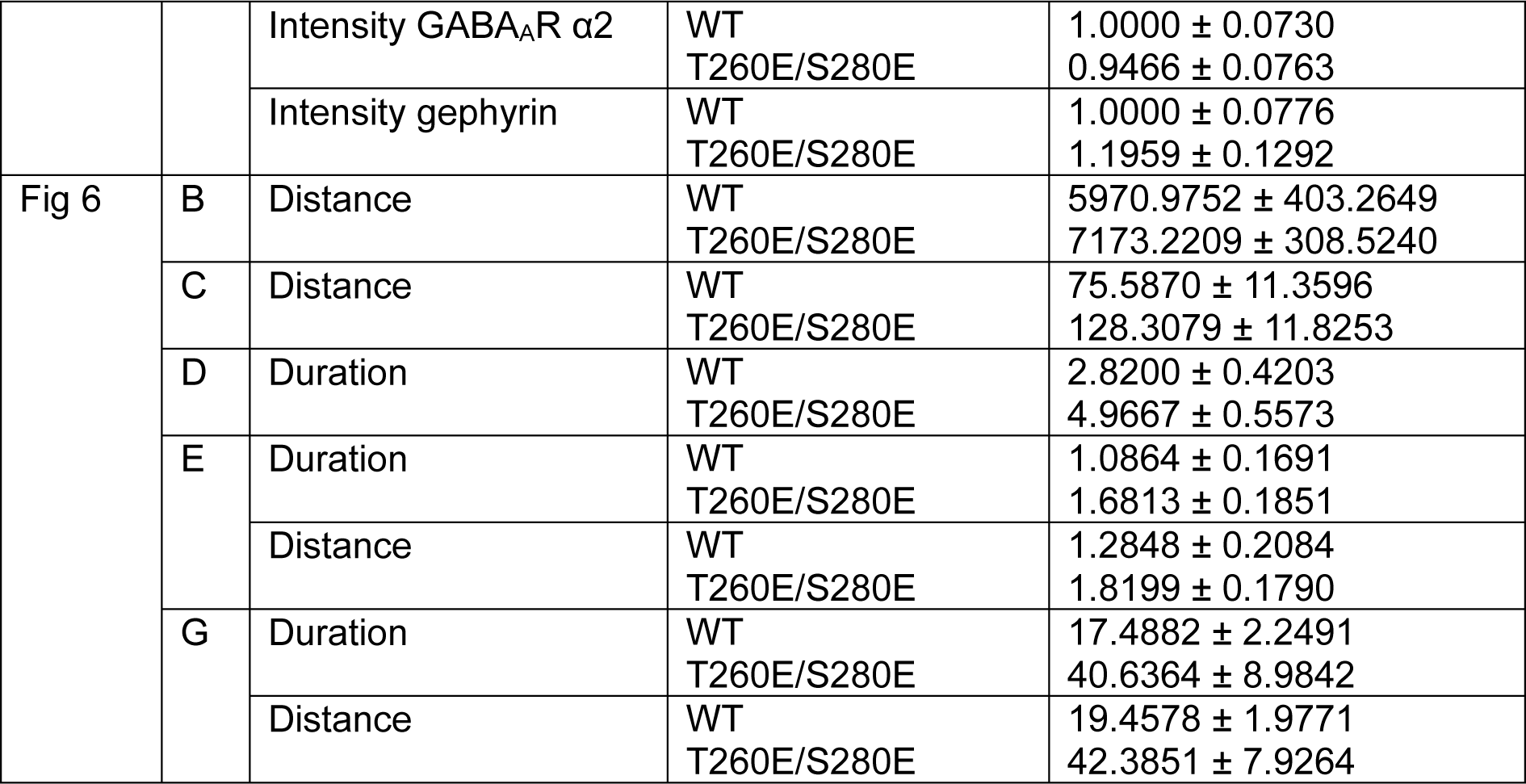
Data summary for principal figures. Summary of median values ± interquartile range (25th–75th percentile) for the Diffusion Coefficient (D) and Explored Area (EA) shown in Fig. 1, and mean values ± SEM for other supplementary data. Related to Figure 1-6.

## Data and code availability

The program used for analysis is available in open access (GitHub - mlrennerfr/Diinamic: Clustering analysis on single-molecule localization microscopy data). The information that supports the development of this protocol is available from the corresponding or technical author upon reasonable request.

The data that support the findings of this study are available from the corresponding author upon reasonable request.

## Acknowledgments

We are grateful for S. Tyagarajan for providing gephyrin-WT-GFP construct. We thank M. Renner for providing MATLAB analysis programs for STORM experiments. We warmly thank E. M. Petrini and J. Dupuis for critical feedback on experiment design. We are also grateful to the Animal Facility of Institut du Fer à Moulin (IFM). This work benefited from the technical support of engineers from the Imaging Facility of IFM (T. Eirinopoulou and M. Savariradjane) and from CNRS UMR 3750 at ESPCI (B. Cinquin). This work was supported in part by Inserm, Sorbonne Université-UPMC as well as by the Agence Nationale de la Recherche (ANR WATT ANR-14-CE13-0032), Fondation pour la Recherche Médicale EQU202203014844, Fondation Française pour la Recherche sur l’Epilepsie and Ligue Française contre l’Epilepsie. STORM/PALM equipment was supported by DIM NeRF from Région Ile-de-France and by the Fondation pour la Recherche sur le Cerveau / Rotary ‘Espoir en tête’.

## Author contributions

S.L and Z.M designed the experiments, prepared the figures and wrote the paper. Z.M. and J.G. performed and analyzed WNK blockade single particle tracking experiments. Z.M. and M.T. performed and analyzed WNK activation single particle tracking experiments. M.T. executed and analyzed surface/total experiments. Z.M. conducted and analyzed STORM experiments, with M.T. assisting for WNK activation experiments. Z.M. performed and analyzed cluster colocalization experiments with R.R. assisting on gephyrin single mutants’ experiments. M.R. prepared the hippocampal cultures. M.R. and E.P. performed calcium imaging experiments and M.R., E.P. and Z.M. analyzed the data. Z.M. conducted stereotaxic viral infection, Z.M. and M.N. verified infection expression. Z.M. and M.N.B. designed and conducted behavior experiments and Z.M analyzed the data. C.D. and C.LM. designed electrophysiological experiments, C.D. performed experiments and analyzed the data. Z.M., M.N. and Z.I. performed immunohistochemistry experiments and analyzed the data.

## Supplemental information

Figure S1 related to Figure 1.

Figure S2 related to Figure 2.

Figure S3 related to Figure 2, 3, 4.

Figure S4 related to Figure 1, 2.

Figure S5 related to Figure 5.

Figure S6 related to Figure 6.

Table S1. Data values related to Figure S1-S6.

