## Supplemental Information for "WNK-Dependent Phosphorylation of Gephyrin Tunes GABA_A_ Receptors at Inhibitory Synapses and Modulates Anxiety Behavior"

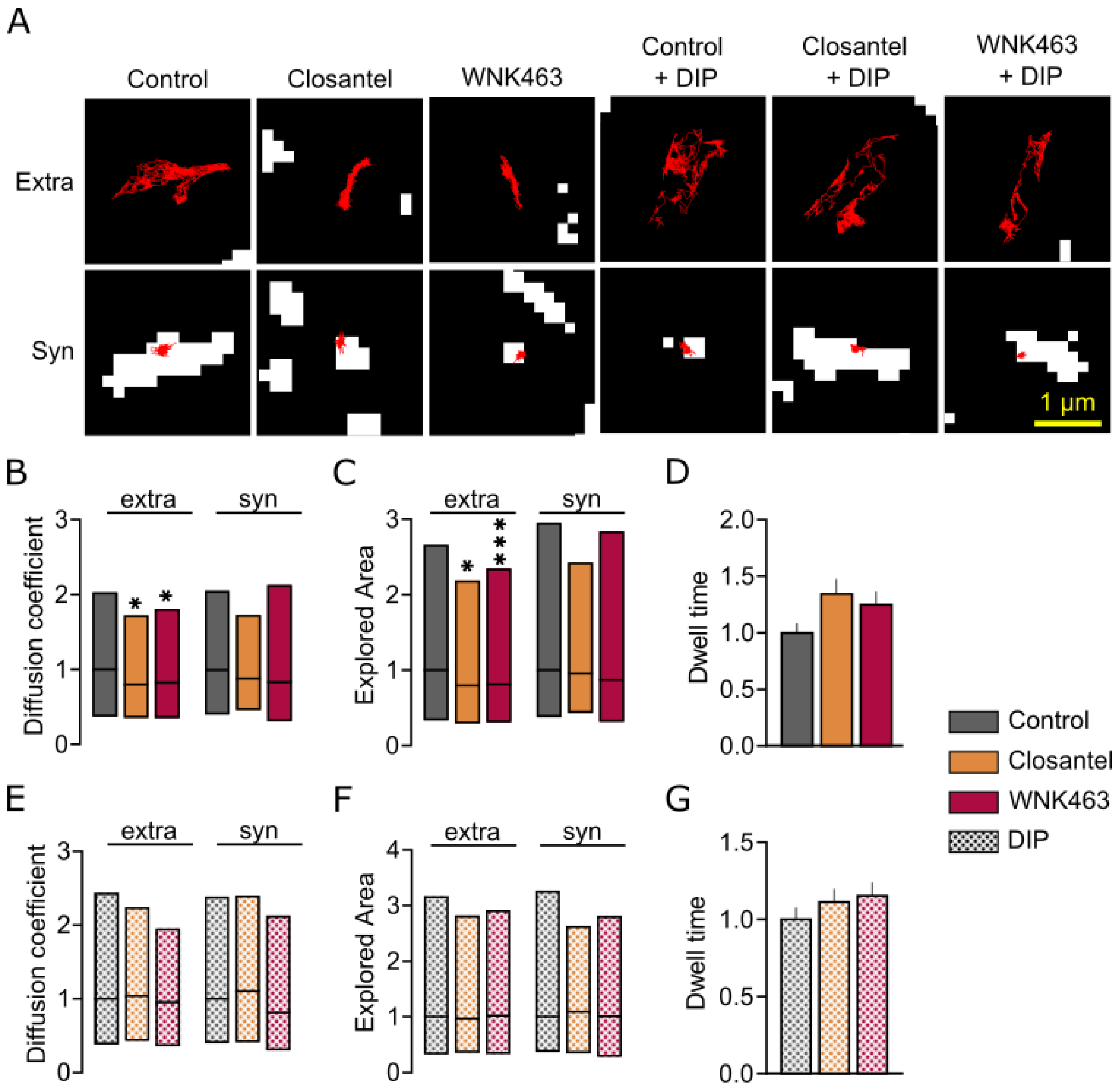


**Figure S1. Acute blockade of WNK signaling confines extrasynaptic GABA$_{A}$R α1 in endocytic wells.**

**A.** Representative images of extrasynaptic (Extra) or synaptic (Syn) GABA$_{A}$R α1 trajectories (red) in neurons maintained in control, Closantel, WNK463, Closantel + dynamin inhibitory peptide (DIP) or WNK463 + DIP conditions. White spots represent gephyrin clusters. Scale bar, 1 µm.
**B, C, E, F.** Diffusion coefficient (B, E) and explored area (C, F) for extrasynaptic (extra) and synaptic (syn) GABA$_{A}$R α1 for neurons treated with closantel (orange), WNK463 (red), closantel + DIP (dotted orange), WNK463 + DIP (dotted red) as compared to their respective controls (grey and dotted grey).***B:*** Control: extra n=905 QDs, syn n=252 QDs, Closantel: extra n=653 QDs, syn n=158 QDs, WNK463: extra n=529 QDs, syn n=163 QDs, 4 cultures. Closantel: extra p=0.0112, syn p=0.7025, WNK463: extra p=0.0312, syn p=0.6853; ***C:*** Control: extra n=2701 QDs, syn n=744 QDs, Closantel: extra n=1954 QDs, syn n=467 QDs, WNK463: extra n=1579 QD, syn n=474 QD, 4 cultures. Closantel: extra p<0.0001, syn p=0.0593,

WNK463: extra p<0.0001, syn p=0.2591; ***E:*** Control + DIP: extra n=563 QDs, syn n=227 QDs, Closantel + DIP: extra n=644 QDs, syn n=224 QDs, WNK463 + DIP: extra n=538 QDs, syn n=202 QDs, 3 cultures. Closantel + DIP: extra p=0.4288, syn p=0.8925, WNK463 + DIP: extra p=0.0711, syn p=0.1823. Data are presented as median values ± 25–75% IQR. Values were normalized and compared to the corresponding control values. Kolmogorov-Smirnov test; ***F:*** Control + DIP: extra n=1686 QDs, syn n=681 QDs, Closantel + DIP: extra n=1932 QDs, syn n=672 QDs, WNK463 + DIP: extra n=993 QDs, syn n=565 QDs, 3 cultures. Closantel + DIP: extra p=0.1138, syn p=0.1395, WNK463 + DIP: extra p=0.4525, syn p=0.0616. Data are presented as median values ± 25–75% IQR. Values were normalized and compared to the corresponding control values. Kolmogorov-Smirnov test.

**D, G.** Dwell time of $GABA_AR$ α1 for neurons treated with closantel (orange), WNK463 (red) (D), closantel + DIP (dotted orange), WNK463 + DIP (dotted red) (G) as compared to their respective controls (grey and dotted grey). ***D:*** Control n=406 QDs, closantel n=245 QDs, WNK463 n=240 QDs, 4 cultures. Closantel p>0.9999, WNK463 p=0.6532; ***G:*** Control + DIP n=342 QDs. Closantel + DIP n=289 QDs, WNK463 + DIP n=305 QDs, 3 cultures. Closantel + DIP p>0.9999, p=0.7880, WNK463 + DIP p=0.8455. Data are presented as mean values ± SEM. Values were normalized to the corresponding control values. Kruskal-Wallis test.

Related to Figure 1.

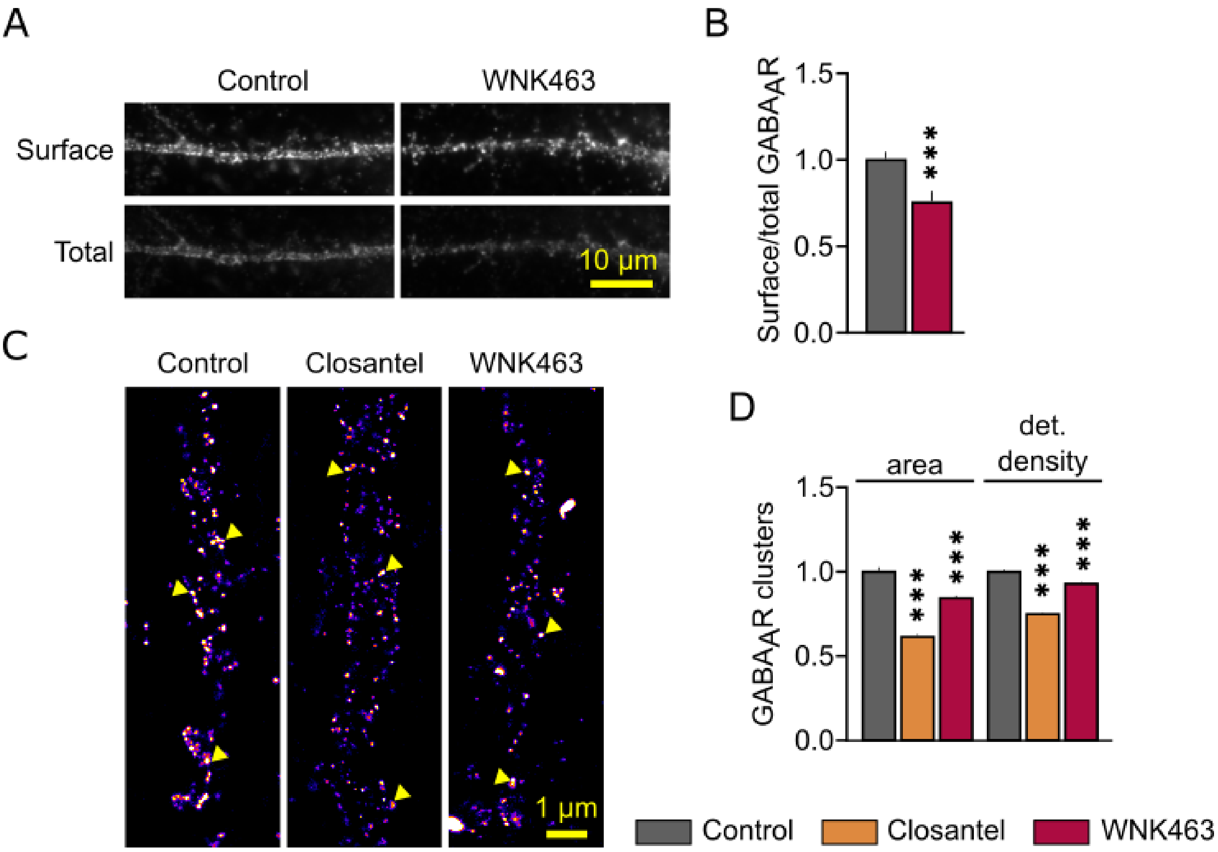


**Figure S2. Reduced GABA$_A$R α1 clustering upon acute blockade of WNK signaling.**

**A.** Conventional fluorescence images of surface GABA$_A$R α1 (top), and total GABA$_A$R α1 (bottom) immunostaining for neurons maintained in control vs WNK463 conditions. Scale bar, 10 µm.

**B.** Surface/total fluorescence intensity ratio of GABA$_A$R α1 for control (grey) vs WNK463 (red) treated neurons. Control n= 40 cells, WNK463= 40 cells, 2 cultures. p=0.0004. Data are presented as mean values ± SEM. Values were normalized and compared to the corresponding control values. Mann-Whitney test.

**C.** STORM rendered images of GABA$_A$R α1 immunostaining for neurons maintained in control, closantel and WNK463 conditions. Warmer colors correspond to higher detection density. Arrowheads show examples of GABA$_A$R α1 clusters. Scale bar, 1 µm.

**D.** Quantification of surface area and detection density of GABA$_A$R α1 clusters of neurons in control (grey) vs closantel (orange) or WNK463 (red) conditions. Control n= 5456 clusters, closantel n=6676 clusters, WNK463 n=7521 clusters. Area: closantel p<0.0001, WNK463 p=0.0010, detection density: closantel p<0.0001, WNK463 p<0.0001. Data are presented as mean values ± SEM. Values were normalized to the corresponding control values. Kruskal-Wallis test.

Related to Figure 2.

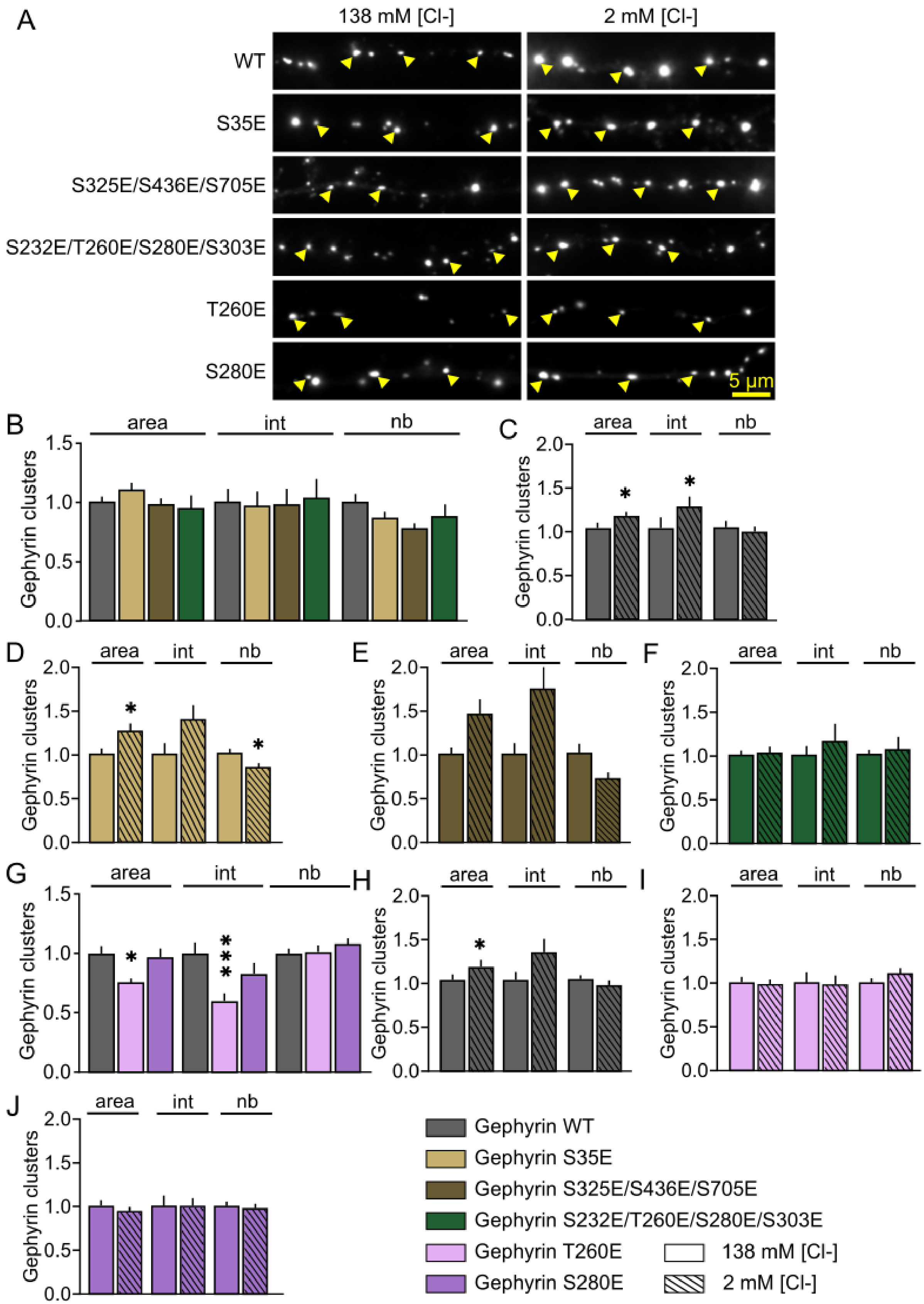
A
138 mM [Cl-]
2 mM [Cl-]
WT
S35E
S325E/S436E/S705E
S232E/T260E/S280E/S303E
T260E
S280E
5 µm
B
C
D
E
F
G
H
I
J
area
int
nb
Gephyrin clusters
Gephyrin WT
Gephyrin S35E
Gephyrin S325E/S436E/S705E
Gephyrin S232E/T260E/S280E/S303E
Gephyrin T260E
Gephyrin S280E
138 mM [Cl-]
2 mM [Cl-]

**Figure S3. Clustering of candidate gephyrin phosphomutants in basal and enhanced WNK activity conditions.**

**A.** Representative images of gephyrin clusters (white) for neurons transfected with WT gephyrin or with phosphomimetic mutants (as indicated) and treated with 138 mM [Cl-] or 2mM [Cl-] solutions. Arrowheads show examples of gephyrin clusters. Scale bar, 5 µm.
**B-F.** Quantification of the mean surface area (area), intensity (int) and number (nb) of gephyrin clusters in neurons expressing gephyrin-WT (grey) or gephyrin-S35E (beige), or gephyrin-S325E/S436E/S705E (brown) or gephyrin-S232E/T260E/S280E/S303E (green) and treated with 138 mM [Cl-] (plain) or 2 mM [Cl-] (hatched) solutions. ***B:*** WT n=37 cells, S35E n=27 cells, S325E/S436E/S705E n=28 cells, S232E/T260E/S280E/S303E, n=30 cells, 3 cultures. Area: S35E p>0.9999, S325E/S436E/S705E p>0.3977, S232E/T260E/S280E/S303E p=0.2878; intensity: S35E p>0.9999, S325E/S436E/S705E p>0.9999, S232E/T260E/S280E/S303E p>0.9999; number: S35E p>0.9999, S325E/S436E/S705E p=0.4132, S232E/T260E/S280E/S303E p=0.3258; ***C:*** 138 mM [Cl-] n=37 cells, 2 mM [Cl-] n=34 cells, 4 cultures. Area p=0.0441; intensity p=0.0493; number p=0.9681. D, 138 mM [Cl-] n=28 cells, 2 mM [Cl-] n=27 cells, 3 cultures. Area p=0.0339; intensity p=0.1200; number p=0.0297. ***E:*** 138 mM [Cl-] n=30 cells, 2 mM [Cl-] n=25 cells, 3 cultures. Area p=0.0765; intensity p=0.2098; number p=0.1032. ***F:*** 138 mM [Cl-] n=27 cells, 2 mM [Cl-] n=23 cells, 3 cultures. Area p=0.6432; intensity p=0.9539; number p=0.4629.
**G-J.** Quantification of the mean surface area (area), intensity (int) and number (nb) of gephyrin clusters in neurons expressing gephyrin-WT (grey) or gephyrin-T260E (pink), or gephyrin-S280E (purple) and treated with 138 mM [Cl-] (plain) or 2 mM [Cl-] (hatched) solutions. ***G:*** WT n=53 cells, T260E n=44 cells, S280E n=53 cells, 7 cultures. Area: T260E p=0.0192, S280E p>0.9999; intensity: T260E p<0,0001, S280E p=0.2076; number: T260E p>0.9999, S280E p>0.9999. Data are presented as mean values ± SEM. Values were normalized to the corresponding control values. Kruskal-Wallis test. ***H:*** 138 mM [Cl-] n=53 cells, 2 mM [Cl-] n=40 cells, 5 cultures. Area p=0.0471; intensity p=0.0736; number p=0.4747. ***I:*** 138 mM [Cl-] n=44 cells, 2 mM [Cl-] n=40 cells, 4 cultures. Area p=0.9751; intensity p=0.1975; number p=0.8411. ***J:*** 138 mM [Cl-] n=53 cells, 2 mM [Cl-] n=49 cells, 5 cultures. Area p=0.5924; intensity p=0.8939; number p=0.9349.
Data are presented as mean values ± SEM. Values were normalized to the corresponding control values. Mann-Whitney test.
Related to Figure 2, 3, 4.

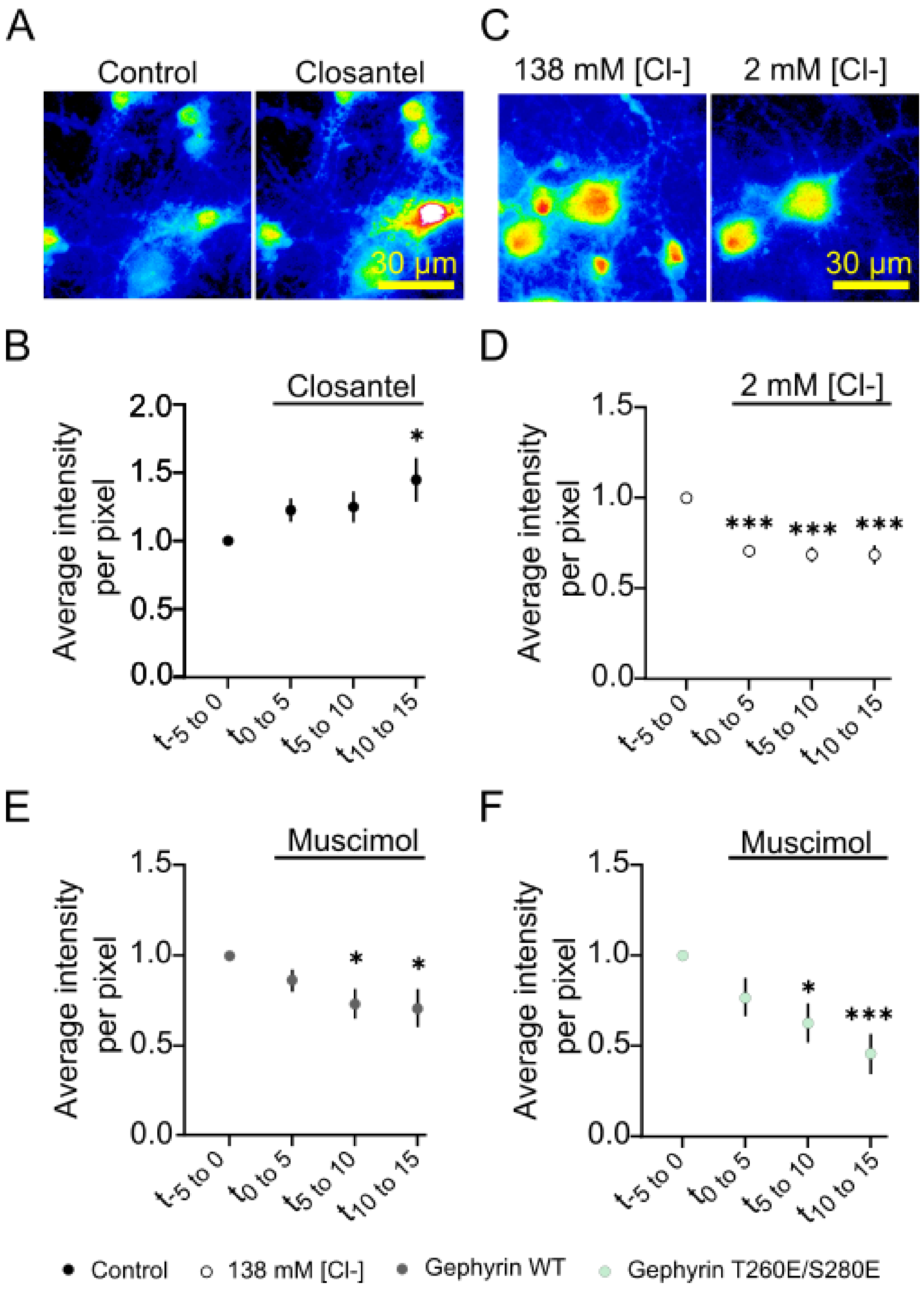
A
Control
Closantel
30 µm
C
138 mM [Cl-]
2 mM [Cl-]
30 µm
B
Closantel
Average intensity per pixel
t-5 to 0
t0 to 5
t5 to 10
t10 to 15
D
2 mM [Cl-]
E
Muscimol
F
Muscimol
Control
138 mM [Cl-]
Gephyrin WT
Gephyrin T260E/S280E

**Figure S4. WNK signaling tunes neuronal activity in hippocampal cultures.**

**A.** Pseudocolor images of neurons loaded with Fluo4-AM before (Control) and after closantel (A) exposure. Warmer colors correspond to higher Fluo4-AM fluorescence intensities. Scale bar, 30 µm.
**B.** Mean Fluo4-AM fluorescence intensity per pixel in region of interest drawn on neuronal somata before and after closantel exposure. n=20 cells, 2 cultures. $t_{0\ \text{to}\ 5}$ p=0.0007, $t_{5\ \text{to}\ 10}$ p=0.0826, $t_{10\ \text{to}\ 15}$ p=0.0266.
**C.** Pseudocolor images of neurons loaded with Fluo4-AM and exposed to 138 mM [Cl-] and then to 2 mM [Cl-] solutions. Warmer colors correspond to higher Fluo4-AM fluorescence intensities. Scale bar, 30 µm.
**D.** Mean Fluo4-AM fluorescence intensity per pixel measured on the soma of neurons exposed successively to 138 mM [Cl-] and 2 mM [Cl-] solutions. n=22 cells, 2 cultures. $t_{0\ \text{to}\ 5}$ p<0.0001, $t_{5\ \text{to}\ 10}$ p<0.0001, $t_{10\ \text{to}\ 15}$ p<0.0001.
**E-F.** Mean Fluo4-AM fluorescence intensity per pixel in region of interest drawn on neuronal somata for Gephyrin WT (F) or Gephyrin T260E/S280E (G) expressing neurons before and after muscimol exposure. ***E:*** n=14 cells, 2 cultures. $t_{0\ \text{to}\ 5}$ p=0.1257, $t_{5\ \text{to}\ 10}$ p=0.0115, $t_{10\ \text{to}\ 15}$ p=0.0424. ***F:*** T260E/S280E n=12 cells, $t_{0\ \text{to}\ 5}$ p=0.1020, $t_{5\ \text{to}\ 10}$ p=0.0136, $t_{10\ \text{to}\ 15}$ p=0.0002.
Data are presented as mean values ± SEM. Values were normalized to the corresponding cells at $t_{-5\ \text{to}\ 0}$. One-Way Repeated Measures ANOVA.
Related to Figure 1, 2.

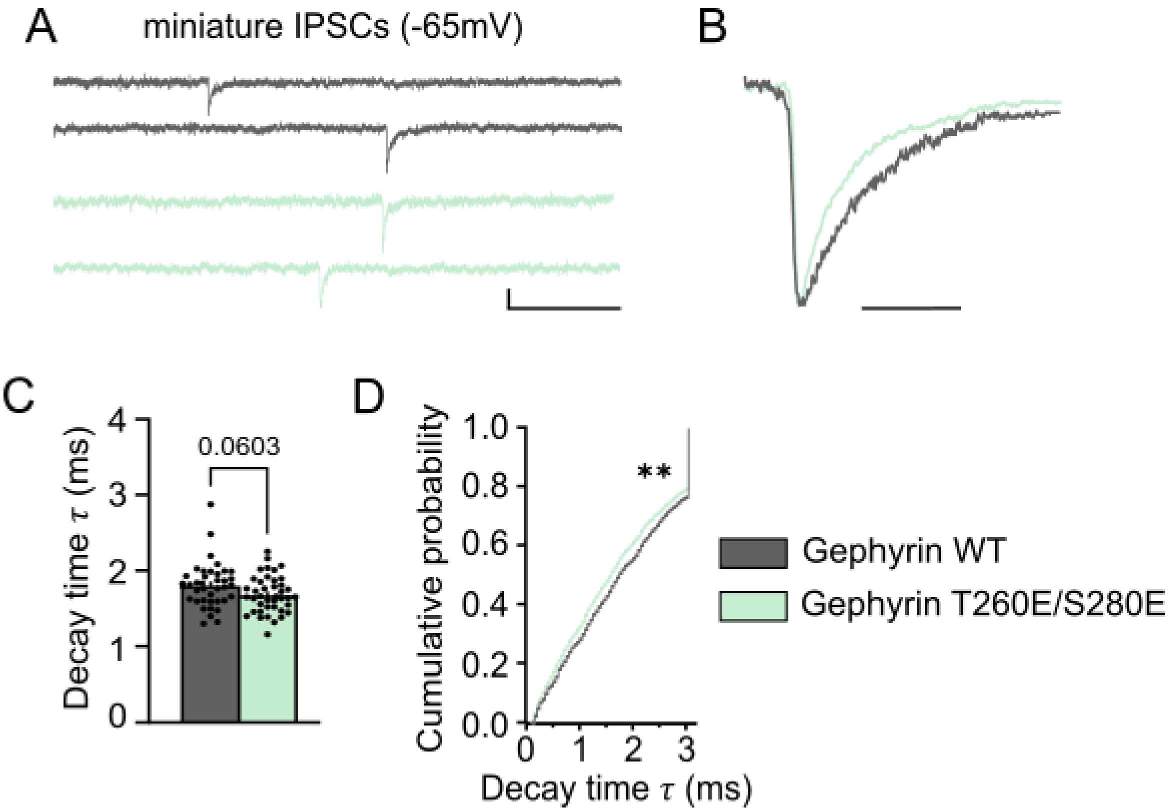


**Figure S5. Hippocampal expression of Gephyrin-T260E/S280E alters GABAergic synaptic current kinetics.**

**A.** Representative recordings of miniature inhibitory postsynaptic currents (mIPSCs) from dentate granular neurons of Gephyrin WT mice (grey) and Gephyrin T260E/S280E infected mice (green). Scale bars, 200 ms, 20 pA.
**B.** Overlay sample of postsynaptic inhibitory events recorded in dentate gyrus neurons from Gephyrin WT (grey) and Gephyrin T260E/S280E infected (green) groups. Scale bar, 10 ms.
**C.** Quantification of mean mIPSCs decay time constant ($\tau$) in Gephyrin WT and Gephyrin T260E/S280E infected cells. WT n=39 cells from 7 animals, T260E/S280E n=41 cells from 7 animals. Data are presented as mean values ± SEM. Mann-Whitney test p=0.0603.
**D.** Cumulative probability of mIPSCs event decay time constant ($\tau$). WT n=1817 events, T260E/S280E n=2282 events, Kolmogorov-Smirnov test p=0.0023.
Related to Figure 5.

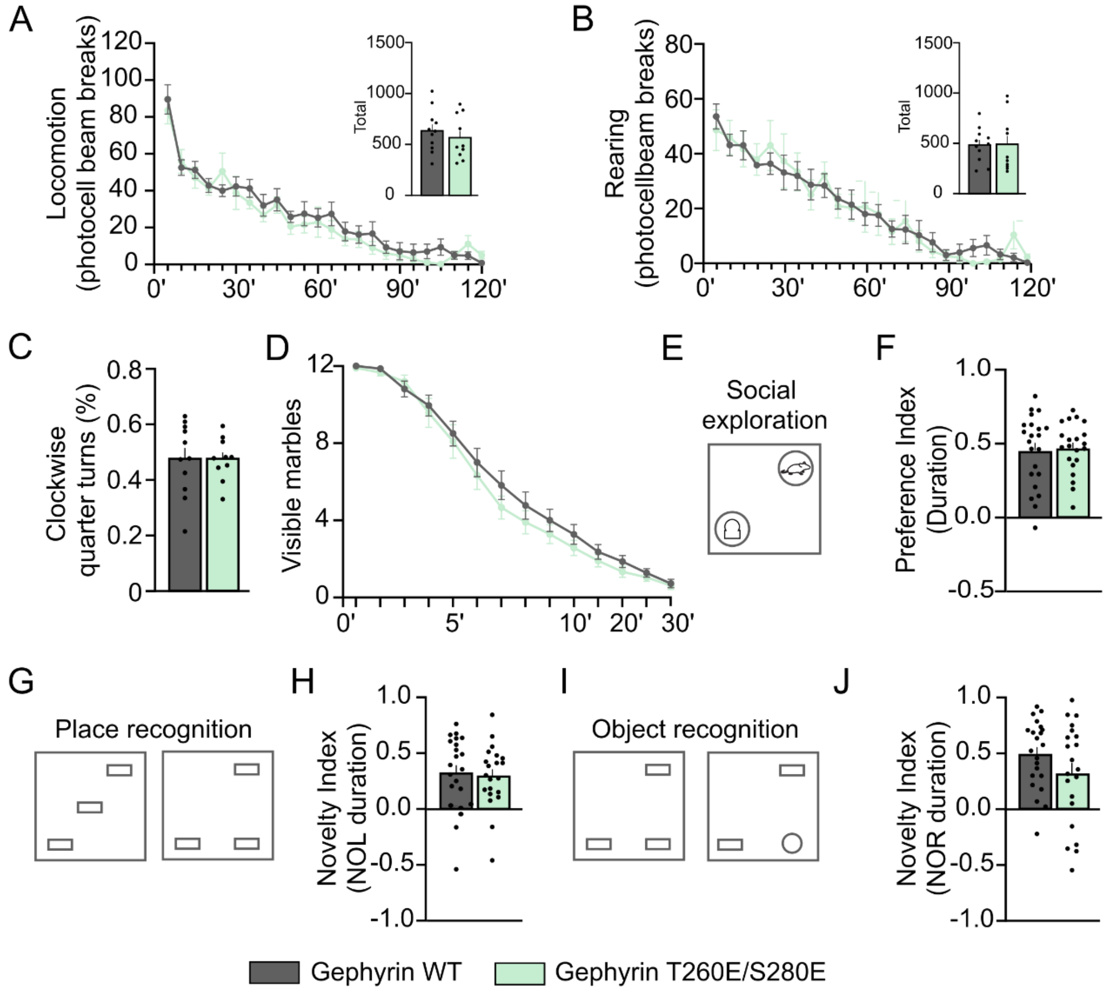


**Figure S6. Hippocampal gephyrin-T260E/S280E overexpression does not affect locomotion, stereotypic-like behavior, social behavior or spatial memory.**

**A-B.** Quantification of locomotion (number of photocell beam breaks) (A) and rearings overtime (B) and total for WT (grey) and T260E/S280E infected mice (light green) in actimeter. WT n=11 animals, T260E/S280E n=10. A. total p=0.4739. B, total p=0.9884.
**C.** Percentage of clockwise quarters turns over total quarter turns in actimeter. WT n=11 animals, T260E/S280E n=10. p=0.9841.
**D.** Quantification of visible marbles over time in marble burying test. WT n=22 animals, T260E/S280E n=21 p=0.4159.
**E.** Scheme representing the social exploration test. Areas containing an unfamiliar object or an unfamiliar mouse are respectively represented on the bottom left and top right of the open-field (OF).
**F.** Preference index for exploration duration. Positive values represent a preference for the unfamiliar mouse. WT n=22 animals, T260E/S280E n=21 animals. p=0.8080.
**G, I.** Scheme of the place recognition test (G), and the object recognition test (I). Left OFs represent old dispositions, right OFs represent the test phases.
**H-J.** Novelty index for exploration duration for the place recognition test (H) and the object recognition test (J). Positive values represent a preference for the new location (H) and the new object (J). WT n=22 animals, T260E/S280E n=21 animals. H, p=0.7722. J, p=0.1435.
Data are presented as mean values ± SEM. t-tests (C, F, H, J) and 2-way ANOVA (D).
Related to Figure 6.

| | | | | |
|---|---|---|---|---|
| Fig S1 | B | Extra D | Control<br>Closantel<br>WNK463 | 1.0000 (0.3726 ; 2.0337)<br>0.8020 (0.3571 ; 1.7225)<br>0.8257 (0.3537 ; 1.8117) |
| | | Syn D | Control<br>Closantel<br>WNK463 | 1.0000 (0.4009 ; 2.0332)<br>0.8809 (0.4602 ; 1.7304)<br>0.8319 (0.3144 ; 2.1054) |
| | C | Extra EA | Control<br>Closantel<br>WNK463 | 1.0000 (0.3358 ; 2.6638)<br>0.79833 (0.2924 ; 2.1859)<br>0.8129 (0.3095 ; 2.3490) |
| | | Syn EA | Control<br>Closantel<br>WNK463 | 1.0000 (0.3783 ; 2.9515)<br>0.9567 (0.4347 ; 2.4141)<br>0.8712 (0.3174 ; 2.8349) |
| | D | Dwell time | Control<br>Closantel<br>WNK463 | 1.0000 ± 0.0839<br>1.3442 ± 0.1339<br>1.2478 ± 0.1203 |
| | E | Extra D | Control +DIP<br>Closantel +DIP<br>WNK463 +DIP | 1.0000 (0.3839 ; 2.4303)<br>1.0396 (0.4295 ; 2.2376)<br>0.9538 (0.3615 ; 1.9457) |
| | | Syn D | Control +DIP<br>Closantel +DIP<br>WNK463 +DIP | 1.0000 (0.4023 ; 2.3589)<br>1.1060 (0.4095 ; 2.3870)<br>0.8143 (0.2982 ; 2.1192) |
| | F | Extra EA | Control +DIP<br>Closantel +DIP<br>WNK463 +DIP | 1.0000 (0.3261 ; 3.1696)<br>0.9690 (0.3535 ; 2.8188)<br>1.0205 (0.3341; 2.9172) |
| | | Syn EA | Control +DIP<br>Closantel +DIP<br>WNK463 +DIP | 1.0000 (0.3697 ; 3.2684)<br>1.0910 (0.346 ; 2.6098)<br>1.0074 (0.2783 ; 2.7818) |
| | G | Dwell time | Control +DIP<br>Closantel +DIP<br>WNK463 +DIP | 1.0000 ± 0.0768<br>1.1128 ± 0.0860<br>1.1539 ± 0.0868 |
| Fig S2 | B | Surface/total Intensity | Control<br>WNK463 | 1.0000 ± 0.0404<br>0.6949 ± 0.0586 |
| | D | Area | Control<br>Closantel<br>WNK463 | 1.0000 ± 0.0195<br>0.6142 ± 0.0074<br>0.8425 ± 0.0095 |
| | | Detection Density | Control<br>Closantel<br>WNK463 | 1.0000 ± 0.0061<br>0.7493 ± 0.0041<br>0.9276 ± 0.0047 |
| Fig S3 | B | Area | WT<br>S35E<br>S325E/S436E/S705E<br>S232E/T260E/S280E/S303E | 1.0000 ± 0.0631<br>0.9879 ± 0.0570<br>0.9546 ± 0.1154<br>1.1118 ± 0.0631 |
| | | Intensity | WT<br>S35E<br>S325E/S436E/S705E<br>S232E/T260E/S280E/S303E | 1.0000 ± 0.1245<br>0.9875 ± 0.1294<br>1.0427 ± 0.1656<br>0.9756 ± 0.1280 |
| | | Number | WT<br>S35E<br>S325E/S436E/S705E<br>S232E/T260E/S280E/S303E | 1.0000 ± 0.0820<br>0.7847 ± 0.0484<br>0.8880 ± 0.1055<br>0.8750 ± 0.0583 |
| | C | Area | 138 mM [Cl-]<br>2 mM [Cl-] | 1.0000 ± 0.0631<br>1.1302 ± 0.0579 |
| | | Intensity | 138 mM [Cl-] | 1.0000 ± 0.1245 |

| | | | | |
|---|---|---|---|---|
| | | | 2 mM [Cl-] | 1.2364 ± 0.1223 |
| | | Number | 138 mM [Cl-]<br>2 mM [Cl-] | 1.0000 ± 0.0820<br>0.9497 ± 0.0674 |
| | D | Area | 138 mM [Cl-]<br>2 mM [Cl-] | 1.0000 ± 0.0580<br>1.2553 ± 0.0883 |
| | | Intensity | 138 mM [Cl-]<br>2 mM [Cl-] | 1.0000 ± 0.1186<br>1.3863 ± 0.1708 |
| | | Number | 138 mM [Cl-]<br>2 mM [Cl-] | 1.0000 ± 0.0527<br>0.8373 ± 0.0541 |
| | E | Area | 138 mM [Cl-]<br>2 mM [Cl-] | 1.0000 ± 0.0714<br>1.4471 ± 0.1733 |
| | | Intensity | 138 mM [Cl-]<br>2 mM [Cl-] | 1.0000 ± 0.1204<br>1.7335 ± 0.3778 |
| | | Number | 138 mM [Cl-]<br>2 mM [Cl-] | 1.0000 ± 0.1141<br>0.7153 ± 0.0719 |
| | F | Area | 138 mM [Cl-]<br>2 mM [Cl-] | 1.0000 ± 0.0525<br>1.0227 ± 0.0757 |
| | | Intensity | 138 mM [Cl-]<br>2 mM [Cl-] | 1.0000 ± 0.1053<br>1.1560 ±0.2055 |
| | | Number | 138 mM [Cl-]<br>2 mM [Cl-] | 1.0000 ± 0.0584<br>1.0596 ± 0.1532 |
| | G | Area | WT<br>T260E<br>S280E | 1.0000 ± 0.0630<br>0.7337 ± 0.0475<br>0.9498 ± 0.0727 |
| | | Intensity | WT<br>T260E<br>S280E | 1.0000 ± 0.0918<br>0.5710 ± 0.0677<br>0.8055 ± 0.0929 |
| | | Number | WT<br>T260E<br>S280E | 1.0000 ± 0.0422<br>0.9934 ± 0.0574<br>1.0569 ±0.0539 |
| | H | Area | 138 mM [Cl-]<br>2 mM [Cl-] | 1.0000 ± 0.0630<br>1.1807 ± 0.0822 |
| | | Intensity | 138 mM [Cl-]<br>2 mM [Cl-] | 1.0000 ± 0.0918<br>1.3549 ± 0.1529 |
| | | Number | 138 mM [Cl-]<br>2 mM [Cl-] | 1.0000 ± 0.0422<br>0.9697 ± 0.0543 |
| | I | Area | 138 mM [Cl-]<br>2 mM [Cl-] | 1.0000 ± 0.0636<br>0.9809 ± 0.0565 |
| | | Intensity | 138 mM [Cl-]<br>2 mM [Cl-] | 1.0000 ± 0.1222<br>0.9787 ± 0.1042 |
| | | Number | 138 mM [Cl-]<br>2 mM [Cl-] | 1.0000 ± 0.0522<br>1.1050 ± 0.0637 |
| | J | Area | 138 mM [Cl-]<br>2 mM [Cl-] | 1.0000 ± 0.0563<br>0.9694 ± 0.0474 |
| | | Intensity | 138 mM [Cl-]<br>2 mM [Cl-] | 1.0000 ± 0.0987<br>1.0286 ± 0.0934 |
| | | Number | 138 mM [Cl-]<br>2 mM [Cl-] | 1.0000 ± 0.0462<br>1.0000 ± 0.0538 |
| Fig S4 | B | $F/F_0$ | $t_{0\ to\ 5}$<br>$t_{5\ to\ 10}$<br>$t_{10\ to\ 15}$ | 1.2240 ± 0.0870<br>1.2484 ± 0.1142<br>1.4471 ± 0.1596 |

| | | | | |
|---|---|---|---|---|
| | D | $F/F_0$ | $t_{0\ to\ 5}$<br>$t_{5\ to\ 10}$<br>$t_{10\ to\ 15}$ | 0.7066 ± 0.0356<br>0.6866 ± 0.0411<br>0.6860 ± 0.0528 |
| | E | $F/F_0$ | $t_{0\ to\ 5}$<br>$t_{5\ to\ 10}$<br>$t_{10\ to\ 15}$ | 0.8643 ± 0.0671<br>0.7313 ± 0.0766<br>0.7086 ± 0.1032 |
| | F | $F/F_0$ | $t_{0\ to\ 5}$<br>$t_{5\ to\ 10}$<br>$t_{10\ to\ 15}$ | 0.7814 ± 0.0911<br>0.6339 ± 0.1032<br>0.4627 ± 0.0883 |
| Fig S5 | C | Decay Time | WT<br>T260E/S280E | 1.8067 ± 0 .0474<br>1.6832 ± 0.0380 |
| Fig S6 | A | Locomotion | WT<br>T260E/S280E | 634.4545 ± 63.0466<br>566.6000 ± 68.3877 |
| | B | Rearing | WT<br>T260E/S280E | 482.7273 ± 51.8235<br>484.2000 ± 88.5910 |
| | C | Turns % | WT<br>T260E/S280E | 0.4714 ± 0.0394<br>0.4705 ± 0.0243 |
| | F | Preference Index | WT<br>T260E/S280E | 0.4627 ± 0.0511<br>0.4783 ± 0.0372 |
| | H | Novelty Index NOL | WT<br>T260E/S280E | 0.3317 ± 0.0705<br>0.3043 ± 0.0616 |
| | J | Novelty Index NOR | WT<br>T260E/S280E | 0.5032 ± 0.0662<br>0.3261 ± 0.0998 |

**Table S1. Data summary for supplementary figures**

Summary of median values ± interquartile range (25th–75th percentile) for the Diffusion Coefficient (D) and Explored Area (EA) shown in Fig. S1, and mean values ± SEM for other supplementary data.

Related to Figure S1-S6.
